# Single Cell Integration Characterises Gastric Cell Metaplasia in Inflammatory Intestinal Diseases

**DOI:** 10.1101/2024.10.11.614415

**Authors:** Amanda J. Oliver, Ni Huang, Ruoyan Li, Raquel Bartolome-Casado, Hogne R. Nilsen, Victoria Gudiño, Elisa Melón-Ardanaz, Michael E. B. FitzPatrick, Nicholas M. Provine, Krzysztof Polanski, Simon Koplev, Lisa Marie Milchsack, Emma Dann, Alexander V. Predeus, Batuhan Cakir, Ken To, Martin Prete, Jonathan A. Chapman, Andrea C. Masi, Emily Stephenson, Justin Engelbert, Sebastian Lobentanzer, Shani Perera, Laura Richardson, Rakesh Kapuge, Anna Wilbrey-Clark, Claudia I. Semprich, Madelyn Moy, Sophie Ellams, Catherine Tudor, Philomeena Joseph, Alba Garrido-Trigo, Ana M. Corraliza, Thomas R.W. Oliver, C. Elizabeth Hook, Julio Saez-Rodriguez, Kylie R. James, Kerstin B. Meyer, Krishnaa T. Mahbubani, Kourosh Saeb-Parsy, Matthias Zilbauer, Marte Lie Høivik, Espen S. Bækkevold, Azucena Salas, Christopher J. Stewart, Janet E. Berrington, Paul Klenerman, Muzlifah Haniffa, Frode L. Jahnsen, Rasa Elmentaite, Sarah A. Teichmann

**Author notes:** Authors contributed equally (Rasa Elmentaite, Sarah A. Teichmann).

## Abstract

The gastrointestinal (GI) tract consists of connected organs, from the oral cavity to rectum, which function to ensure efficient nutrient uptake and barrier immunity. Diseases of the GI tract affect millions worldwide and as such there are now over 25 published single cell RNA-sequencing (scRNAseq) datasets surveying the GI tract, profiling specific anatomical regions, cell lineages, ages and diseases. To consolidate these efforts, we harmonised and integrated scRNAseq datasets across the whole GI tract from developing and adult human tissues, as well as newly generated data from preterm gut. We uniformly processed 385 samples from 189 healthy controls using a newly developed automated QC approach (scAutoQC). In total, our healthy reference contains ∼1.1 million cells which we annotated to a total of 137 fine-grained cell states. We anchor 13 published and 1 unpublished GI disease datasets covering gastric and colorectal (CRC) cancers, celiac disease, ulcerative colitis (UC) and Crohn’s disease (CD) to this reference, taking our atlas to a total of 1.6 million cells. We provide our atlas as a valuable resource to the community (available at gutcellatlas.org). Using this resource, we discover epithelial cell metaplasia arising from stem cells across intestinal inflammatory diseases (celiac, UC and CD) and CRC with transcriptional similarity to cells of the gastric and Brunner’s glands. Whilst previously linked to mucosal healing, we now implicate these cells in inflammation through recruitment of immune cells including T cells and neutrophils, and through direct interactions with T cells. Overall, we discover a shift in paradigm whereby changes in stem cells during inflammation lead to altered mucosal tissue architecture, which in turn contributes to ongoing inflammation. These findings highlight that in addition to barrier function, epithelial cells actively contribute to progression of inflammation which may be a function applicable to other tissues and diseases.

## Introduction

The human gastrointestinal (GI) tract is a complex system comprising several organs that work together to efficiently absorb nutrients, while simultaneously providing an immunologically active barrier. Diseases of the GI tract are prevalent; ulcerative colitis (UC) and Crohn’s disease (CD) affect over 7 million people worldwide, and 2 million new colorectal cancer (CRC) cases are diagnosed annually^1,2^. Recently, the field of single cell transcriptomics has revolutionised our understanding of GI health, development, and disease, offering unprecedented molecular insights^3–7^. Over 25 single-cell RNA sequencing (scRNAseq) studies of the human GI tract have been published to date, with most primarily focused on profiling individual organs or specific upper (e.g., oesophagus, stomach) and lower (e.g., small and large intestine) GI regions. The integration of these publicly available datasets not only serves as a valuable resource for the Human Cell Atlas community and beyond^8^, but also enables seamless cross-regional comparisons of intestinal cell types.

The epithelial cells that line the lumen of the GI tract develop from a common endoderm progenitor and acquire their regional identity early in embryogenesis^9^. In some cases, this regional identity can be altered in adulthood and lead to metaplasia, where mature tissue is replaced by cells normally occurring in other anatomical regions^10^. Cross-regional comparison from single cell data is especially relevant for metaplasia, enabling the direct comparison between healthy cells and their metaplastic counterparts. Intestinal metaplasia is well described in the stomach and in Barrett’s oesophagus patients, where the mucosa is transformed to intestinal epithelial cells, and is associated with increased risk of gastric and esophageal adenocarcinomas^11,12^. However, stomach-like metaplasia of intestinal cells is less well characterised. These lineages have been assigned various names including pyloric metaplasia, pseudopyloric metaplasia, gastric metaplasia, ulcer-associated cell lineage (UACL), and spasmolytic polypeptide-expressing metaplasia (SPEM), and mostly studied using histology^10,13–15^. The prevailing hypothesis is that these metaplastic lineages may be part of the mucosal healing process in the epithelial layer and may also transition to neoplastic cells^10^. However, the origin and functional role of metaplastic cells in both acute and chronic tissue damage remains unresolved.

In this study, we assemble a healthy reference GI tract atlas, carefully curating and integrating 23 publicly available scRNAseq datasets from both developing and adult human GI tissues, as well as 2 unpublished datasets from preterm and adult gut. Using this atlas as a healthy reference, we anchor 13 publically available and 1 unpublished GI disease datasets to resolve cell type features across disease sets. In total, our atlas integrates and harmonises data from 271 donors across health, development and disease, resulting in 1.6 million cells which we annotate to 137 fine-grained cell types. We provide this atlas as a resource to the community, and use it to interrogate cell types and signatures in inflammatory intestinal diseases.

Analysing cell types in scRNAseq disease data across the GI tract, we discover the presence of two distinct populations representing pyloric-type metaplastic cells in the terminal ileum of pediatric and adult inflammatory bowel disease (IBD) patients and the duodenum of celiac disease patients. Using our reference to deconvolute bulk-RNA-sequencing data, we identify the presence of these cells across multiple studies of IBD and CRC. Here, we show that these metaplastic lineages arise from stem cells in inflamed tissues, and retain stem-like characteristics. These cells have transcriptional features resembling *MUC6*+ mucous gland neck cells and *MUC5AC*+ surface foveolar cell types of the healthy stomach, and to some extent duodenal Brunner’s gland cells. However, in inflammatory intestinal diseases we find that *MUC6*-expressing cells can be distinguished by expression of inflammatory programs such as upregulated MHC (major histocompatibility complex) class II and immune cell-recruiting chemokines. Our in-depth analysis suggests that this metaplastic cell type, which we termed INFLAREs (<u>Infla</u>mmatory <u>e</u>pithelial cell<u>s</u>), interact with CD4+ T cells and recruit monocytes, dendritic cells and neutrophils to the site of damage *via* interaction with activated vascular endothelial cells. Thus, we propose a shift in epithelial stem cells which alters the differentiation pathway from healthy to metaplastic lineages and in turn contributes to ongoing inflammation in chronic disease settings.

## Results

### Pan-gastrointestinal data integration

We curated, integrated and harmonised healthy cells across the GI tract profiled by scRNA-seq data from 23 published and 2 unpublished datasets (Figure 1a-c, Supplementary Table 1, Extended data 1a, b). Tissues covered in the atlas broadly include the oral mucosa, oesophagus, stomach, small and large intestines and mesenteric lymph nodes. We harmonised metadata from each study including age, sex, fine-grained anatomical region, sampling methods such as sequencing technology, cell and tissue fraction enrichment (Extended data 1c, Supplementary Table 2). To reduce batch effects from different genome references, sequence alignment and quality control (QC) methods, we remapped raw data and processed gene counts uniformly through a newly developed automated QC pipeline (Methods, Figure 1b, Extended data 1 and 2). Our pipeline (scAutoQC) removes the reliance on manual thresholds with low quality cells/droplets identified by taking into account 8 standard metrics (such as mitochondrial and ribosomal genes) in a reduced QC metric space. Cells/droplets are filtered out if present within a cluster that fails QC according to thresholds set by Gaussian Mixture modelling (Extended data 2a). A major advantage of our approach is the calculation of thresholds using multiple QC metrics per cell, exploiting both the distribution of individual metrics as well as correlations between them, allowing retention of cells with unique features such as plasma cells which have lower numbers of expressed genes. After initial filtering, we removed 596,449 (31.22%) low quality cells which were present within failed QC clusters and cells from samples where < 10% of cells passed QC. Next, we removed a further 67,846 cells predicted as doublets by scrublet, with further manual doublet removal after integration and cell type annotation. To integrate data, we used scVI^16^, which has been shown to perform well in single cell data integration benchmarking studies^17^.

**Figure 1:**
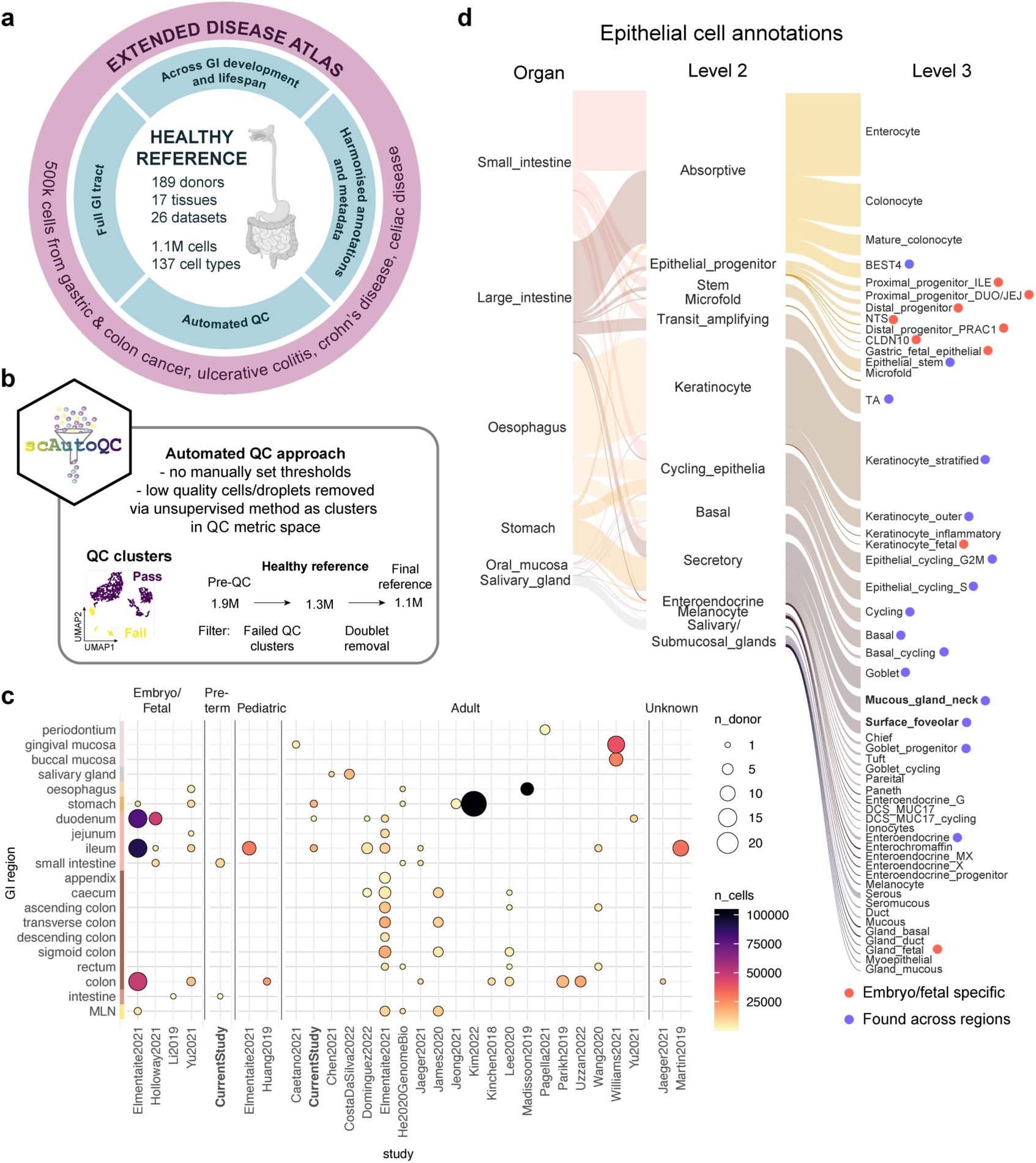
Overview of pan-gastrointestinal cell integration. a) Schematic overview of the atlas denoting the healthy reference as a core, encompassing developing and adult full GI tract, with additional disease datasets mapped onto the healthy reference by transfer learning. b) Overview of scAutoQC, an automated QC pipeline which uses an automated, unsupervised approach to remove low quality cells/droplets within the QC metric space. c) Overview of the number of cells and donors per study (x axis), broken down by age and region (y axis) of the GI. Dot size indicates the number of donors, colour indicates the number of cells. Colour bar on y axis indicates broad level organs (oral mucosa, salivary gland, oesophagus, stomach, small intestine, large intestine and MLN, mesenteric lymph node). d) Sankey plot overviewing epithelial cell annotations at fine-grained level and organ level. Cell types appearing only in development and appearing across multiple regions are highlighted. Cell types in bold are a major focus of the paper. Link thickness indicates the proportion of cells within each category. Link colour relates to the organ (left side) and level 2 annotation (right side).

The final integrated data was annotated into 6 broad lineages which were subclustered and further annotated into fine-grained cell types (Supplementary Figure 1-3). Due to large heterogeneity in our dataset in terms of GI regions and life stages (Extended data 3a, b), we further subclustered epithelial and mesenchymal cells, which had the most heterogeneity by age and/or region, to accurately annotate fine-grained cell types (Extended data 1). Cell types were annotated in a semi-automated method, with manual annotations based on known marker genes cross-referenced with automated annotation from multiple CellTypist^18^ models of published studies^3,4,19,20^. In total, we arrived at a healthy reference atlas comprising ∼1.1 million cells from 156 adult/pediatric and 33 embryonic/fetal/preterm donors, which we annotated to 137 fine-grained cell types (Figure 1d, Extended data 1). We annotated 51 epithelial cell types/states highlighting specialised epithelial populations during development and across the GI tract, along with common populations found across regions (Figure 1d). We found particularly high granularity in immune cell populations with 17 T/NK, 16 myeloid, and 11 B cell and plasma cell subsets.

Altogether, we used our reference atlas to dissect similarities and differences in cell types across GI regions and life stages. In particular, we focused on cells that can be grouped into three categories. Firstly, unappreciated cell types that have been found in one region, but whose comprehensive distribution along the GI was not fully appreciated – for example, BEST4 cells in stomach (Supplementary Figure 2d), *MUC6*+ mucous gland neck (MGN) and *MUC5AC+* surface foveolar cells in the duodenum (Supplementary Figure 2e). Secondly, rare cell types that are less than 0.02% of cells in the atlas, such as deep crypt secretory (DCS) cells in the colon (Supplementary Figure 2f), which were also identified in a recent integration study^21^, and Langerhans dendritic cells (DCs) (Supplementary Figure 1a). Finally, the integrated atlas allowed increased distinction between highly similar cell types such as IgA1 and IgA2 B plasma cells (Supplementary Figure 1d).

### Cellular changes in the healthy GI tract

Comparing cell type composition in developing versus mature (adult and pediatric) stomach, duodenum, ileum and colon, we observed broad enrichment of neural and mesenchymal cells in developing tissues (Extended data 3c). Myeloid populations, especially macrophages (*CD209*+ and *LYVE1*+), were generally enriched in developing compared to adult intestinal tissues, but not in developing vs adult stomach (Extended data 3c, d). In line with the co-development of intestinal IgA responses and microbiota after birth^22^, we found enrichment of most B cell subsets in mature GI (Extended data 3d). An exception was progenitor B cells, which were enriched in all developing GI tissues, as previously seen in fetal intestines^19^ (Extended data 3d). Although most T cell populations were enriched in mature adult GI tissues, there was enrichment of ILC3 and CD56^bright^ cytotoxic NK cells in developing GI (Extended data 3d).

Differential abundance across GI regions in adult/pediatric tissues revealed enrichment of endothelial cells in oral mucosa compared to other regions (Extended data 3e), which is consistent with this region being highly vascularised^23^. At fine-grained cell annotation, we observed enrichment of IgA2 and IgM plasma cells in the oesophagus (Extended data 3f). In mesenchymal populations, we found a number of region-specific fibroblasts enriched in oral mucosa, oesophagus and rectum (Supplementary Figure 1a, Extended 3g).

In summary, our integration of healthy tissues presents the most comprehensive cell reference encompassing lineages along the full GI tract in humans and allows interrogation of the shared and unique cell lineages along the GI tract.

### Disease-relevant cell changes across IBD types

After defining the healthy reference annotations, we integrated additional data from 63 disease donors and 5 diseases, including UC, CD, pediatric IBD, CRC and gastric cancers. Similar to our approach in the healthy dataset, we started from the raw data, remapped and applied the scAutoQC approach to the disease data. This ensured that the healthy and disease references are comparable. To annotate the disease data, we first used the healthy reference to predict broad lineages using scANVI and scArches.

We found an apparent reduction in epithelial cell numbers in disease compared to healthy datasets, which could be due to epithelial damage and/or focus on immune cells in many studies of disease. Since we were specifically interested in identifying metaplastic cells, we added new epithelial data by incorporating count matrices from a large recently published study of CD^24^ and unpublished data from celiac disease. In total we added ∼500,000 cells to our atlas, bringing the total number to 1.6 million cells across 27 studies and 271 donors. To annotate cells at fine-grained resolution, we used scANVI and scArches, projecting disease data onto our subclustered, lineage-and region-specific views of the atlas (Methods, Extended data 1b and 4). Overall, we successfully annotated disease cells using this approach, with the exception of a population of gastric cancer cells with high uncertainty metric which are likely cancer cells. We also observed a distinct cluster of cells in large intestinal epithelium that were originally labelled as a mixture of goblet cells and doublets (Extended data 4c, e-j). Upon closer inspection, these cells were reannotated as Paneth cells based on marker genes *DEFA5, DEFA6, REG3A* and *PLA2G2A* (Figure 2c, Extended data 4g, j). Paneth cell metaplasia is a known feature of chronic inflammation of the colon^25,26^. Indeed, Paneth cells were found in multiple donors with IBD, including in neighbouring inflamed tissue, but not in the healthy colon (Figure 2c, Extended data 4h).

**Figure 2:**
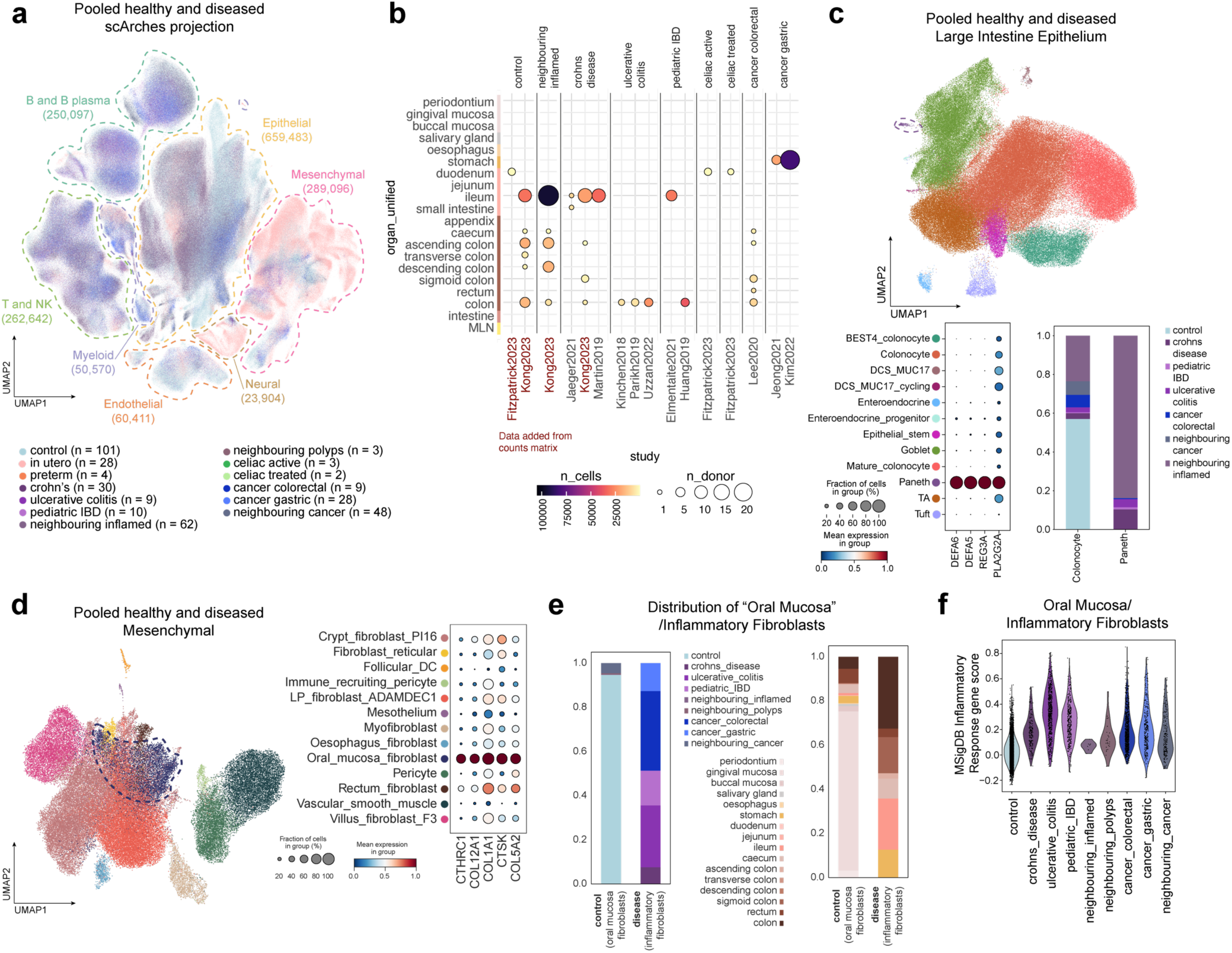
Projection of disease data highlights metaplastic cell lineages in IBD. a) UMAP of joint healthy and disease atlas with cells coloured by disease category with n = number of donors per category. Dotted lines indicate broad cell lineages with cell numbers indicated in brackets. b) Overview of the extended disease data showing number of cells and donors per study, broken down by disease. Dot size indicates the number of donors, colour indicates the number of cells. Studies with red text (Fitzpatrick and Kong) were not remapped, instead data was added to the atlas as count matrices. c) UMAP of large intestine epithelial cells from adult/pediatric healthy and disease samples, highlighting metaplastic Paneth cells. Dotplot of Paneth cell markers and proportions of cells from control and disease categories of colonocytes vs Paneth cells which are not present in healthy large intestine. d) UMAP of mesenchymal populations from healthy and diseased adult/pediatric tissue, with “oral mucosa fibroblasts” highlighted with marker gene dotplot. d) Cell proportions of “oral mucosa fibroblasts”/inflammatory fibroblasts in control and disease across GI regions in the healthy reference only and pooled healthy and disease atlas. Oral mucosa fibroblasts are highlighted as a cell type present in the intestines only in disease. e) Comparison of oral mucosa fibroblasts from healthy samples (reference) and disease samples (query) showing increased expression of inflammatory/activated fibroblast markers from literature ^27^. f) Violin plot of MSigDB Inflammatory Response gene score in oral mucosa fibroblasts across disease categories. Pathway is significant from gene set enrichment analysis comparing differentially expressed genes between oral mucosa fibroblasts in healthy vs diseased samples (Extended data 5).

Transfer learning of annotations for the fibroblast compartment on disease samples revealed a population of fibroblasts from IBD and cancer labelled as oral mucosa fibroblasts, despite coming from the stomach, small and large intestine (Figure 2d, e, Extended data 5a-c). In the healthy reference, this population included cells from gingival mucosa, with marker genes involved in collagen and matrix deposition (*CTHRC1*, *COL12A1*, *COL1A1*, *COL5A2, CSTK*). Compared with healthy counterparts, disease-associated oral mucosa-like fibroblasts overexpressed marker genes of inflammatory/activated fibroblasts reported in UC and CD^27^. Moreover, using CellTypist models from published studies^3,4^, oral mucosa fibroblasts were predicted as inflammatory/activated fibroblast populations. Hence we refer to this cell population from disease samples as inflammatory fibroblasts. Hierarchical clustering of oral mucosa/Inflammatory fibroblasts across locations distinguished cells from gingiva/periodontium but not buccal mucosa, potentially reflecting unique microbiome and disease susceptibilities of gingival mucosa^28,29^. Furthermore, in disease-associated inflammatory fibroblasts vs healthy oral mucosa fibroblasts, DEGs were enriched for various inflammatory pathways (KEGG IL-17 signalling pathway, MSigDB Hallmark Interferon gamma response, MSigDB Hallmark inflammatory response), supporting an immune-mediated role of this population in the lower GI tract during disease. In periodontitis, gingival mucosa fibroblasts also upregulate inflammatory genes, particularly those involved in recruiting neutrophils (*CXCL1, 2, 5 and 8*)^30^. This oral mucosa fibroblast state, which is homeostatic in healthy gingival mucosa, may be primed to promote inflammation and resolve infection. Gingival mucosa is particularly vulnerable to damage and infection, as the first point of contact for commensals and pathogens entering the GI tract, and due to injury exposure through mastication. Thus, the existence of a readily primed fibroblast state in homeostasis may facilitate rapid infection and healing responses. In the intestines, this population does not exist in homeostasis and only arises in severe inflammatory environments, perhaps similar to inflammation of the gingival mucosa. Understanding the regulation of inflammatory gene expression programs in this fibroblast population, especially in gingival mucosa, could give clues for novel therapeutic strategies to target fibroblasts in IBD.

To identify populations that are consistently enriched in disease, we applied differential abundance method Milo on integrated healthy and disease datasets^31^. Importantly, we used both disease and matching healthy controls overlayed on the atlas reference to compute the differences in disease samples. This approach has been recently shown to lead to a very precise identification of disease-enriched cell states using single cell data^32^. We observed oral mucosa/inflammatory fibroblasts enriched in CD, along with monocytes, plasmablasts, IgG plasma cells, *TREM2*+ macrophages and migratory DCs (Extended data 6a).

### Epithelial metaplasia in intestinal disease

In the epithelial compartment of the small intestine, we observed two distinct populations of cells with unique expression signatures across both healthy and diseased samples. In the healthy duodenum, we observed *MUC6*-expressing mucous gland neck (MGN) cells and *MUC5AC*-expressing surface foveolar cells which resembled cells of the Brunner’s glands of the duodenum (Figure 3a, b, Supplementary Figure 2e) ^33,34^. Intriguingly, we also identified similar populations enriched in the ileum of patients with IBD (Figure 3c, Extended data 6a, b). In addition, we observed more *MUC6*-expressing cells in the duodenum of patients with untreated celiac disease (Figure 3c, Extended data 6b).

**Figure 3:**
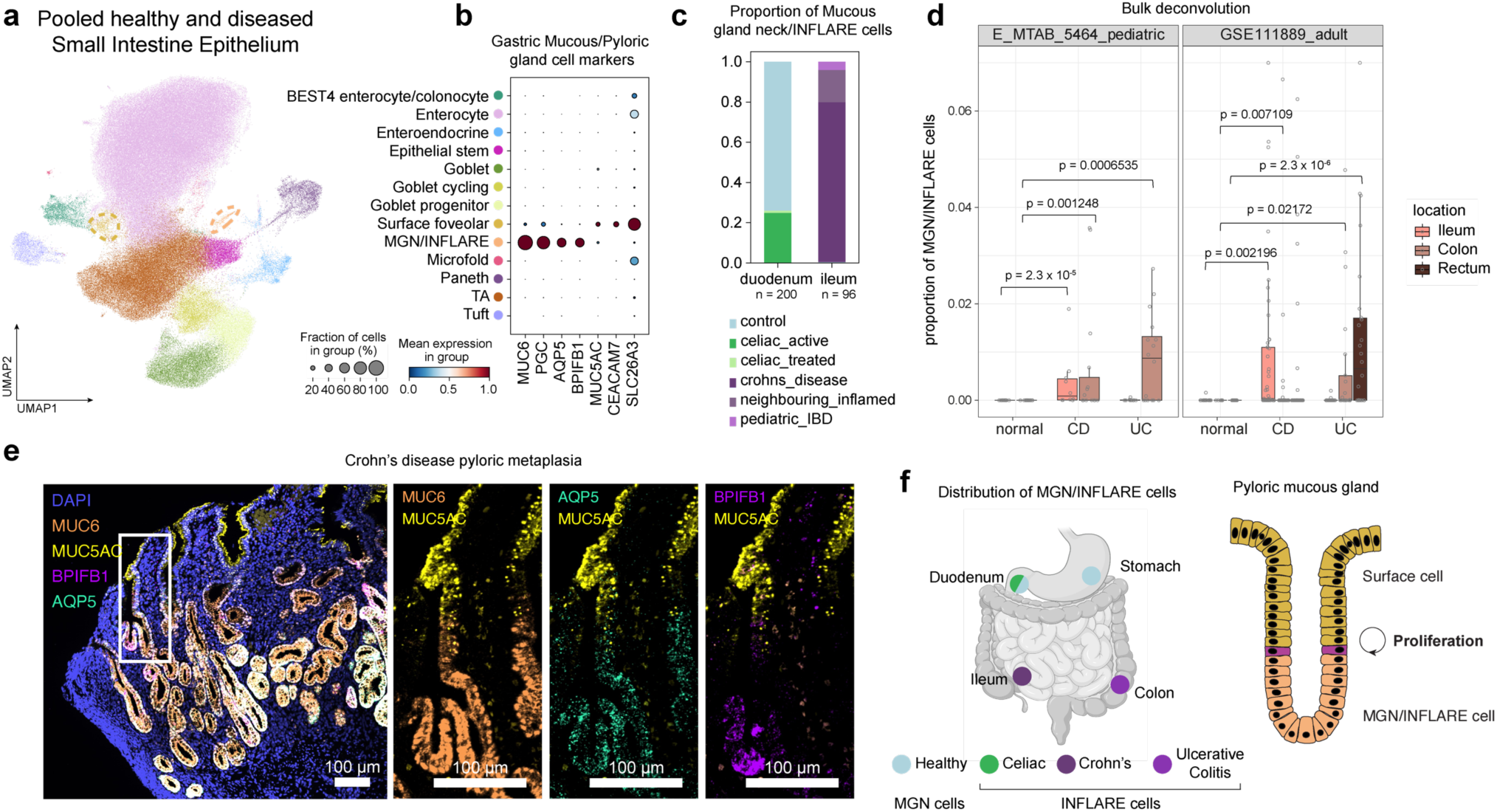
Integration across GI identifies INFLAREs - <u>Infla</u>mmato<u>ry</u> <u>E</u>pithelial cells resembling pyloric mucous gland/Brunner’s gland neck cells in healthy stomach and duodenum. a) UMAP showing cells from small intestinal epithelium in the full atlas (healthy and diseased). MGN (mucous gland neck)/INFLARE and surface foveolar cells, both involved in pyloric mucous gland metaplasia, are highlighted with a dashed circle. b) Marker gene dot plot of gastric mucous/pyloric gland cell markers (MGN and surface Foveolar cells). c) Proportion of MGN/INFLARE cells by disease category in duodenum and ileum. d) Bulk deconvolution (BayesPrism) using disease intestinal epithelium as a reference in studies of crohn’s disease (CD) and ulcerative colitis (UC). For E_MTAB_5464 n = 25 (CD), n = 27 (UC) and n = 27 (normal). For GSE111889 n = 122 (CD), n = 71 (UC) and n = 50 (normal). The lower edge, upper edge and centre of the box represent the 25th (Q1) percentile, 75th (Q3) percentile and the median, respectively. The interquartile range (IQR) is Q3 - Q1. Outliers are values beyond the whiskers (upper, Q3 + 1.5 x IQR; lower, Q1 - 1.5 x IQR). e) smFISH staining of MGN/INFLARE cell marker genes (*MUC6, AQP5, BPIFB1*) and surface foveolar cell markers (*MUC5AC*) in duodenum biopsy from a patient with CD and pyloric metaplasia. f) Schematic of MGN/INFLARE cell distribution across the stomach and intestines and structure of pyloric mucous glands/Brunner’s glands.

Marker genes of the *MUC6*-expressing population included *MUC6*, *PGC*, *AQP5* and *BPIFB1* and of the *MUC5AC*-expressing population included *MUC5AC*, *CEACAM7* and *SLC26A3* (Figure 3b). Within the *MUC5AC*-expressing cluster, we observed heterogeneous expression including enhanced expression of *CEACAM7*, *CEACAM1*, *DUOX2* and *LCN2* in IBD (Extended data 6c). Intriguingly, we observed significant differences between *MUC6*-expressing cells in the healthy stomach and duodenum (mucous gland neck, MGN, cells) compared with diseased *MUC6*-expressing cells in celiac duodenum and IBD ileum. From herein, we refer to this population in disease as <u>Infla</u>mmatory <u>E</u>pithelial cell<u>s</u> (INFLAREs). We hypothesise that together, these are metaplastic epithelial cell lineages that have been described previously as gastric mucous/pyloric gland metaplasia by histological analysis^13^. Having identified these metaplastic lineages in single cell data for the first time, we are now able to investigate their precise role(s) at a molecular level in disease.

Histology-based gastric metaplasia has previously been reported in approximately 40% of patients with IBD (average from 9 published studies across 729 tissue sections)^13,35–42^. In our atlas, we find INFLAREs in only a small number of patients, which could be reflective of factors such as sampling biases and disease severity. To generalise our findings, we investigated publicly available bulk RNA-seq datasets from three independent cohorts of mucosal biopsies from both pediatric and adult IBD patients. Deconvoluting these datasets using BayesPrism^43^ and our atlas as a reference, we found significantly higher proportions of INFLAREs in CD and UC tissue than in healthy tissue (Figure 3d, Extended data 6d). Reassuringly, INFLAREs were present in both small and large intestine in CD but only in the large intestine in UC patients, which is consistent with the aetiology and site of inflammation in the two conditions (Figure 3d). In addition, we performed deconvolution on TCGA data from colon adenocarcinomas and bulk RNAseq datasets from celiac disease. In colon adenocarcinoma, INFLARE proportions were elevated compared to healthy controls, and particularly elevated in microsatellite instability (MSI)-high cancers, which display higher mutational burden and increased infiltration of immune cells^44^ (Extended data 6e). In duodenal samples from celiac disease, proportions of INFLAREs were higher, albeit not significantly, when compared to healthy duodenum (Extended data 6f).

To further validate the presence of INFLAREs in patients with intestinal inflammatory disease (IBD and celiac), we performed both immuno-histochemistry (IHC) and multiplexed smFISH in a number of patients from different disease cohorts (Supplementary Table 3). Using both techniques, we located INFLAREs (MUC6+*AQP5*+*BPIFB1*+) at the crypt base and metaplastic surface foveolar cells (MUC5AC+) at the crypt top in CD samples (Figure 3e, Extended data 6g, h). Similar to metaplastic surface foveolar cells, we noted heterogeneity in INFLAREs based on co-expression of the marker genes *AQP5* and *BPIFB1* (Extended data 6i). Consistent with previous reports, we found INFLAREs in close association with ulcerated regions and tertiary lymphoid structures in some tissue sections (Extended data 6g). We also found INFLAREs in disease tissue from untreated celiac and UC patients (Extended data 6j, k). In untreated celiac patients, we observed MUC6+ cell localisation within the epithelial layer rather than the submucosa (Extended data 6j, upper panel), demonstrating that INFLAREs are distinct from Brunner’s glands (Extended data 6j, lower panel).

In summary, we describe gastric mucous gland metaplasia, consisting of two distinct populations, at single cell resolution for the first time. We observe heterogenous *MUC5AC*+ cells, representing metaplastic cells equivalent to foveolar cells at the surface of gastric glands. Importantly, we highlight *MUC6*+ INFLAREs in untreated celiac duodenum, CD ileum and UC colon, and epithelial cells resembling MGN cells in the healthy stomach and duodenum (Figure 3f). As in healthy gastric mucous/Brunner’s glands, MUC6+ cells reside at the base of these glands and express mucins, digestive enzymes and pH regulating genes.

### Origin of INFLARE cells

To interrogate the origin of INFLAREs, we performed trajectory analysis (Monocle3) on small intestine epithelial cells. We found that MGN/INFLAREs were branching from the *LGR5+* stem cell population (Figure 4a) and retained a number of stem-like genes along the trajectory (Figure 4b). Using smFISH, we found *LGR5* and *MKI67* expression in INFLAREs in tissue from Crohn’s disease ileum (Figure 4c). We identified a shift in gene expression in *LGR5+* stem cells from IBD patients compared to healthy controls with significantly increased expression of stem cell marker genes (*REG1A, OLFM4, SLC12A2)* (Extended data 7a). We observed enhanced expression in IBD stem cells of interferon stimulated genes such as *IFI27*, MHC class II antigen presentation genes (*HLA-DRA, HLA-DRB1*) and the transcription factor STAT1, which is implicated in interferon induced MHC-II expression. Intriguingly, MHC-II expression in murine *Lgr5+* intestinal stem cells is important for epithelial cell remodelling during infection, through direct interaction with CD4+ T helper cells^45^. In addition, STAT1 has recently been implicated in regulating hematopoietic stem cell maintenance and self-renewal^46^. Thus, stem cells in inflamed tissue differ from those in healthy tissue, and have the potential to give rise to metaplastic cells.

**Figure 4:**
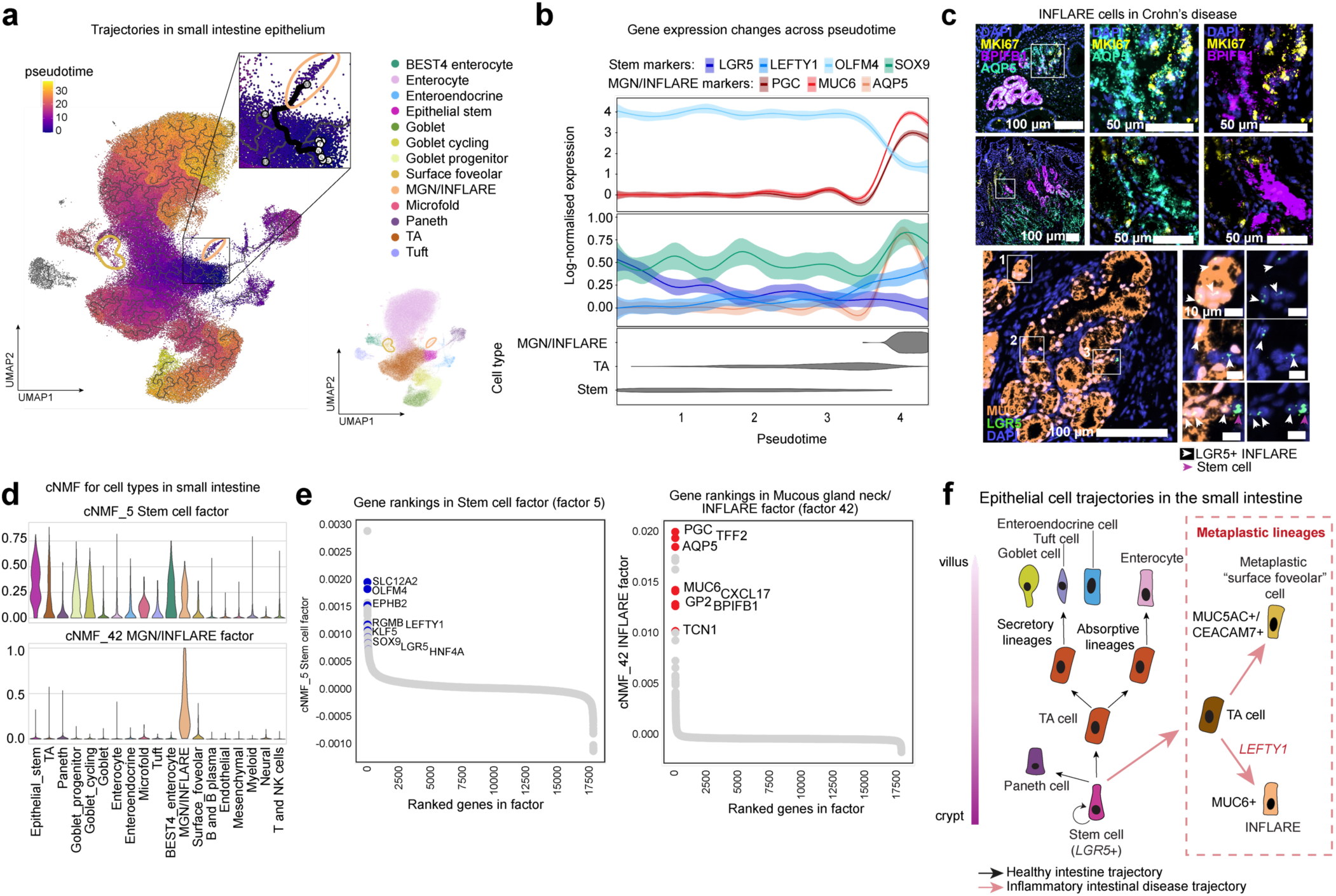
INFLAREs originate from stem cells and retain stem-like properties. a) UMAP of small intestine epithelial cells coloured by pseudotime values inferred with Monocle3 analysis on epithelial cells from ileum only, highlighting trajectory from stem cells to MGN/INFLAREs. Small UMAP alongside indicates cell types. b) Expression of Stem and MGN/INFLARE markers along pseudotime, along with cell type abundance along the Stem - TA - MGN/INFLARE trajectory. c) Proliferation (*MKI67*) and Stem (*LGR5*) marker expression by smFISH in INFLAREs (*MUC6*) from CD ileum and duodenum. d) Non-negative matrix factorisation analysis (Methods) of cell types from small intestine in the atlas. Violin plots showing expression of ranked genes in factors related to MGN (mucous gland neck)/INFLAREs (<u>Infla</u>mmatory <u>E</u>pithelial cell<u>s</u>) and stem cells (*LGR5*+). e) Gene rankings of genes in factors 5 (Stem cell factor) and 42 (MGN/INFLARE factor). Highlighted genes are involved in stem-cell function (blue) and MGN/INFLARE marker genes (red). f) Schematic of epithelial cell trajectories along the crypt-villus axis in the small intestine known during health (black arrows) and hypothesised in our study in inflammatory intestinal diseases (red dotted box and red arrows).

Using unbiased consensus non-negative matrix factorisation (cNMF, Methods), we further distinguished transcriptional programs in common between *LGR5+* stem cells and INFLAREs, supporting their stem-like properties. We also identified factors distinguishing MGN/INFLAREs from other mucous secreting cells, such as surface foveolar and goblet cells (Figure 4d, e, Extended data 7b). In the factor distinguishing goblet cells from INFLAREs, we identified goblet-specific genes (eg. *MUC2*) which are absent from INFLAREs. At the same time, there are shared genes expressed in both goblet cells and INFLAREs, including some commonly expressed genes across mucous-secreting cells reflecting the mucus-secreting function of INFLAREs (Extended data 7c, d). In the factors distinguishing surface foveolar cells, we found a number of mucin encoding genes and *CEACAM* family members, along with genes involved in ion transport and IgA responses (Extended data 7e). Overall, NMF analysis confirms that INFLARE cells are a distinct cell type and have unique features that are not shared with goblet cells or metaplastic surface foveolar cells.

The factor in common across *LGR5+* stem cells and MGN/INFLAREs also connected other less mature cell types (eg. TA cells and Goblet progenitors), and included high ranking expression of many intestinal stem cell marker genes including *SLC12A2, RGMB,* and *LGR5*. *LEFTY1,* another high-ranking gene in this factor, has recently been shown to mark altered undifferentiated stem cell populations that give rise to intestinal metaplasia in the oesophagus (Barrett’s oesophagus)^12^ and in stomach^47^. In the stomach, this population is a small subcluster of MGN cells in healthy stomach mucosa, referred to as “linking” stem cells expressing *LEFTY1* and *OLFM4*. In mammary gland epithelial cells, LEFTY1 was shown to regulate self renewal and drive proliferation in breast tumorigenesis^48^. In our atlas, we found *LEFTY1* expression in INFLAREs and epithelial stem/progenitor populations with highest expression in inflamed conditions (Extended data 7f, g). Together, our data suggests that metaplasia arises from changes within crypt-based stem cells in inflamed tissue giving rise to metaplastic surface foveolar cells and INFLAREs (Figure 4f). We find that INFLAREs retain stem-like properties in intestinal disease, representing a plastic population.

### Metaplasia plays a dual role in mucosal healing and inflammation

Previous studies suggest that metaplasia is an injury adaptation in mucosal tissues in response to wound closure and healing, given the antimicrobial and mucin production capacity of these lineages^10,49^. Indeed, we found that metaplastic surface foveolar cells from ileum in IBD patients had increased expression of antimicrobial related genes compared with healthy counterparts in the stomach and duodenum (Extended data 8a). Additionally, both surface foveolar cells and INFLAREs expressed the trefoil factor *TFF3,* which is normally expressed by goblet cells and has a key role in mucosal healing^50^. This expression profile is in contrast to healthy surface foveolar and MGN cells in the stomach and duodenum which expressed *TFF1* and *TFF2*, respectively (Extended data 8a). In addition, we found that metaplastic surface foveolar cells in IBD ileum expressed high levels of *PIGR*, the polymeric immunoglobulin receptor, and *CCL28* (Extended data 8b) a chemokine crucial for the recruitment of immune cells, in particular IgA plasma cells. Thus, surface foveolar cells in inflamed ileum can enhance recruitment of IgA plasma cells and transcytosis of IgA antibodies, strengthening the mucosal barrier. Overall, our data supports a role of gastric mucous/pyloric gland metaplasia in mucosal healing, with a major contribution from metaplastic surface foveolar cells.

Despite this role for metaplasia in mucosal healing, we found specifically that INFLAREs rather than metaplastic surface foveolar cells, had additional characteristics that may contribute to ongoing chronic intestinal inflammation. We compared MGN/INFLAREs from across different tissues, life stages and diseases in our atlas and found distinct features of MGN/INFLAREs in different tissues and in development (Extended data 8c). Comparing marker genes of MGN/INFLAREs, we found greater similarity between diseased INFLAREs and healthy MGN cells in the stomach than in healthy duodenum (Extended data 8d). Comparing differentially expressed genes between MGN/INFLAREs in our atlas, we found upregulation of inflammatory pathways such as response to cytokines and IFNγ mediated signalling in INFLAREs compared to MGN cells in healthy duodenum and stomach, similar to our comparison of ileal stem cells from CD vs healthy individuals (Extended data 8e-f).

To further interrogate the result of inflammatory signalling from INFLAREs in disease, we performed cell-cell interaction analysis (CellPhoneDB and CellChat, Methods). We found that INFLAREs overexpressed a number of chemokines, for example *CXCL16*, predicted to interact with *CXCR6* on various T cell subsets (Figure 5a, b, Extended data 8g). INFLAREs across tissues and healthy MGN cells in the stomach expressed higher levels of chemokines than MGN cells in healthy duodenum (Figure 5a, Extended data 8g). In addition, INFLAREs expressed much higher levels of chemokines such as *CXCL2*, *3*, *5* and *17* than metaplastic surface foveolar cells (Extended data 8g-i). The INFLARE specific chemokine *CXCL17*, is known to be expressed in mucosal barrier tissues^51^ (Extended data 8h). Although the receptor for CXCL17 is unknown, it is a potent chemoattractant for DCs, monocytes and macrophages and plays a role in angiogenesis^52,53^. In addition, *CXCL2/3/5* neutrophil recruiting chemokines expressed on INFLAREs were predicted in our data to interact with *ACKR1*, an atypical chemokine receptor which can transport chemokines into the vessel lumen ^54^. *ACKR1* has been described as a marker of activated endothelial cells and as part of a cellular module associated with resistance to anti-TNF and anti-integrin α4β7 in IBD^4,55^. Interestingly, a recent study has shown that interactions between vessels and neutrophils can upregulate *ACKR1*^56^. Using smFISH on CD ileum tissue, we found a close association of *ACKR1* vessels with INFLAREs (*MUC6+*) at the base of metaplastic glands (Figure 5c, Extended data 8j). We find that venous endothelial cells (EC_venous) are consistently correlated with INFLAREs in deconvoluted bulk RNAseq datasets of CD, in agreement with staining in tissue (Extended data 8k). Although we did not capture neutrophils in the atlas, we find that neutrophil marker genes (*CXCR1, CXCR2, FCGR3B, PROK2*) are correlated with INFLAREs in bulk RNAseq datasets (Extended data 8k). In summary, INFLAREs express elevated levels of chemokines, with potential to recruit neutrophils and T cells into inflamed tissue during intestinal inflammatory diseases.

**Figure 5:**
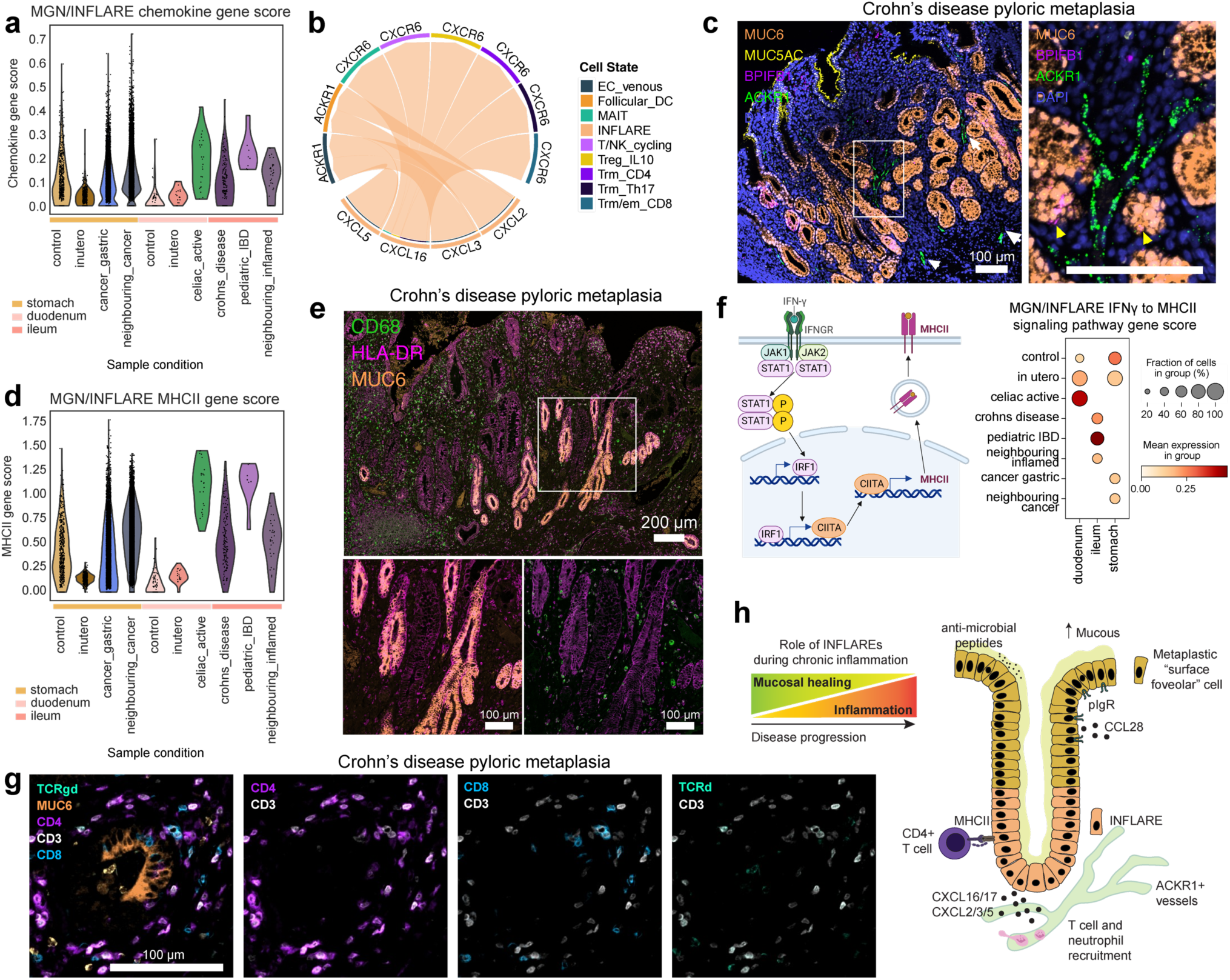
INFLAREs recruit and interact with immune cells in IBD. a) Gene score of chemokines across MGN (Mucous gland neck)/INFLAREs (<u>Infla</u>mmatory <u>E</u>pithelial cell<u>s</u>) from stomach, duodenum and ileum across different conditions. b) Cell-cell interactions mediated by *CXCL* chemokines expressed by INFLAREs and various immune cells/venous endothelial cells. c) smFISH staining of INFLARE (*MUC6, BPIFB1*), Surface foveolar (*MUC5AC*) and activated endothelial cells (*ACKR1*) showing proximity of vessels to metaplastic glands in CD. d) Gene score of MHC-II genes and peptide processing genes across MGN/INFLARE cells from stomach, duodenum and ileum across different conditions. e) Protein staining of INFLAREs (MUC6), macrophages (CD68) and MHC-II (HLA-DR) in ileum from CD resection showing high MHC-II expression in INFLAREs. f) Schematic of signalling pathway from *IFNGR* to *MHC-II* with dotplot of gene scores from this pathway in MGN/INFLAREs from stomach, duodenum and ileum across different conditions. g) Protein staining of INFLAREs (MUC6) and CD4 T cells (CD3+CD4+), CD8 T cells (CD8+CD3+) and γδ T cells (TCRγδ+CD3+) showing interaction between CD4 T cells and INFLAREs. h) Schematic of the potential role of gastric mucous/pyloric gland metaplasia in inflammatory intestinal diseases. INFLAREs and metaplastic surface foveolar cells arise in response to local inflammation in order to promote mucosal healing. As disease progresses INFLAREs contribute to ongoing inflammation through association with activated vessels, the recruitment of various immune cells and direct interactions with CD4 T cells via MHC-II.

In addition to inflammatory cytokines, we found that INFLAREs expressed significantly higher levels of MHC-II related genes compared with MGN cells in healthy tissue (Figure 5d, Extended data 8e-g). In particular, MHC-II related pathways were upregulated specifically when comparing INFLAREs in ileum to MGN cells in healthy duodenum, but not in healthy stomach (Extended data 8e-f). We confirmed the high level of MHC-II protein expression (HLA-DR) in ileum tissue sections from CD patients. INFLAREs (MUC6+ cells) had much higher HLA-DR expression than surrounding MUC6-glands and surface epithelium (Figure 5e, Extended data 8l). Expression of MHC genes is known to increase in response to IFNγ, which is prominent in intestinal inflammation ^57,58^. While some elevation in MHC-II expression was seen in other epithelial cell types in disease samples, including metaplastic surface foveolar and *LGR5+* stem cells, this was markedly increased in INFLAREs compared with MGN cells in healthy tissue (Extended data 8g). In support of the role of IFNγ signalling, we observed increased gene expression in IFNγ signalling and the IFNγ receptor to MHC-II pathways in INFLAREs from inflamed tissue compared with neighbouring inflamed and healthy tissues, which correlated with the levels of MHC-II (Figure 5d, f, Extended data 8e, g, m). In addition, we observed various T cell subsets (CD8, CD4, γδ T cells) surrounding INFLAREs in our IHC imaging of CD and celiac disease tissue (Figure 5g, Extended data 8l, n, o). Consistent with expression of MHC-II, we observed close interaction between CD4+ T cells and INFLAREs (MUC6+) in CD ileum, suggesting that INFLAREs may act as non-conventional professional antigen presenting cells in chronic inflammatory settings.

In summary, by integrating RNAseq data from 1.6 million single cells across the GI tract, we identified metaplastic cell populations in inflammatory intestinal diseases, which resemble gastric mucous glands. Although these cells were rare in single cell data, gastric mucous/pyloric gland metaplasia is estimated to be present in ∼40% of IBD patients. We hypothesised that these metaplastic lineages arise due to inflammatory shifts in intestinal stem cells, leading to altered cell trajectories and transformation of healthy tissue. In addition to the previously hypothesised role in mucosal healing we find that metaplastic cells, specifically INFLAREs, may additionally contribute to ongoing inflammation through recruitment and interaction with T cells and neutrophils (Figure 5h).

## Discussion

Cells across the GI tract play a vital role in nutrient uptake and immunity, and diseases affecting this system have a significant global burden. Despite the prevalence of GI diseases, there is still much to uncover about the shared and tissue specific cell types in health and the cellular and molecular changes in GI disease. To address this gap in knowledge, we curated and integrated single cell datasets covering the whole human GI tract. In contrast with other recent integration efforts^21,59^, we uniformly processed raw data through our bioinformatics workflow including automated QC (scAutoQC), which is of use to future single cell studies and integration efforts. This approach removed processing-related batch effects and uncovered rare cell types that might otherwise be excluded by conventional QC approaches. In total, we integrated 1.6 million cells from 271 donors and 27 studies across health, development and disease which we provide as a resource to the community and interrogate in order to further understand IBD in cellular and molecular detail.

We identified regional differences in health and disease and highlighted various metaplastic cell lineages that lost their regional identity in chronic disease. In the large intestine of IBD patients, we identified Paneth cells, a known metaplastic lineage in colonic inflammatory settings. In our healthy atlas, we identified multiple tissue-specific fibroblast populations, including fibroblasts specific to oral mucosa which had transcriptional similarity to inflammatory fibroblast populations previously characterised in IBD. Therefore, metaplasia may not be confined to epithelial cell types but may occur in other lineages such as fibroblasts. Inflammatory fibroblasts are associated with ongoing inflammation, tissue damage and therapeutic resistance in IBD patients^4,60,61^. We propose that oral mucosa fibroblasts exist in a primed cell state which can be readily activated in response to inflammation such as the inflammatory fibroblast populations in gingival mucosa during periodontitis^30^. Understanding this process may give insights into how intestinal fibroblasts transition to an inflammatory state in IBD and importantly, how this can be halted or reversed.

We identified two distinct metaplastic lineages expressing *MUC6* and *MUC5AC* at single cell resolution in inflammatory intestinal disease, resembling cells in healthy stomach mucous glands and duodenal Brunner’s glands. In disease, we termed *MUC6* expressing cells equivalent to mucous gland neck (MGN) cells as INFLAREs (<u>Infla</u>mmatory <u>E</u>pithelial cell<u>s)</u>. The presence of these cells is best described in the healthy human stomach, and have recently been described in a study using high resolution single cell and spatial methods in healthy duodenal Brunner’s glands^34^. Metaplastic “UACL” glands have been profiled at a transcriptional level by laser capture microdissection, highlighting many of the marker genes identified in our study along with the expression of LGR5 and MKI67 in metaplastic glands^15^. At single cell resolution, a study of pediatric treatment-naive CD and functional gastrointestinal disorder patients identified *MUC6+TFF2+* and *BPIFB1*+*AQP5+* populations, albeit annotated as goblet cells^62^. Similarly another recent single cell study of CD and UC identified MGN/INFLAREs, annotated as *MUC6*+*PGC+DOUX2+* enterocytes, enriched in inflamed ileum of CD^63^. Integration of 12 single cell studies investigating IBD also revealed *DUOX2*+ epithelial cells in UC, which co-express *LCN2* and may be similar to metaplastic surface foveolar cells reported here^21^. Aberrant mucin expression and gastric mucous/pyloric gland metaplasia in CD patients has been reported extensively from histology^13^ and is emerging in transcriptional studies. Here, we identify and interrogate gastric mucous/pyloric gland metaplasia at the single cell level, with full transcriptional resolution. Importantly, we highlight distinguishing features of metaplastic cells from their healthy counterparts in the stomach and duodenum and define epithelial transformation both in stem cells and mature, differentiated cells across intestinal inflammatory diseases.

Our observations suggest a paradigm shift, where metaplasia arises due to a change in adult stem cell identity and potential. Recent studies investigating intestinal metaplasia in oesophagus^12^ and stomach^47^, propose that metaplastic lineages emerge from altered undifferentiated stem cells. In the ileum of IBD patients, we propose a similar transformation following mucosal injury in the small intestine, whereby the intestinal stem cells transform to LEFTY1+ stem cells and differentiate to stomach-like gland cells. We provide multiple orthogonal lines of evidence for stem cell and proliferative features in INFLAREs using cNMF, trajectory analysis and imaging.

Although difficult to dissect the exact origins of altered stem cell dynamics, it is possible that inflammatory signalling in *LGR5*+ stem cells drives intestinal tissue transformation. Inflammatory driven alterations have been identified in stem cells across tissues including neural and hematopoietic stem cells^64–66^. Differentiation trajectories of intestinal organoids from mice are skewed by T helper responses (Th1, 2, 17) where, for example, Th1 responses increased Paneth and goblet cell differentiation^45^. During sustained inflammation, stem cells may be unable to re-enter homeostasis, and differentiation could be further skewed towards metaplastic lineages. Indeed, *ex vivo* colonic organoids cultured from UC patients suggest permanent changes in stem cells with increased expression of anti-microbial and gastric associated genes compared to healthy controls^67^. In a murine model of graft versus host disease, *Lgr5*+ stem cells undergo epigenetic reprogramming with hyper-accessibility in genomic regions associated with interferon signalling and MHC-II antigen processing and presentation^68^. Thus, epithelial transformation such as gastric mucous/pyloric gland metaplasia, may be a direct consequence of lasting changes within stem cells and their trajectories in response to chronic inflammation.

Previous observations suggest that gastric mucous gland metaplasia plays a role in epithelial wound healing and barrier repair after mucosal injury^10^. Our results build on these observations and propose a role of INFLAREs in chemokine-mediated immune cell recruitment and antigen presentation. Increased expression of MHC-II on intestinal epithelial cells in IBD patients has been previously described, along with the ability of epithelial cells to stimulate CD4+ T cells in an MHC-II dependent manner^69,70^. We propose that INFLAREs recruit and similarly interact directly with CD4+ T cells under inflammatory conditions. In addition, INFLAREs can recruit neutrophils, similar to inflammatory fibroblasts^61^, using a cellular circuit likely aided by the close association with *ACKR1*+ vessels found in our data. In support of a disease promoting role, many genes expressed by INFLAREs have been implicated in GWAS studies of IBD, including chemokines *CXCL1/2/3/5* and *JAK2/IFNGR2/STAT1*, involved in IFNγ signalling^71^. Overall we hypothesise that in addition to mucosal healing, the alteration in mucosal tissue structure can in turn sustain inflammatory processes, contributing to the development and maintenance of chronic symptoms in patients with IBD.

Intriguingly, there is a potential link between INFLAREs and CRC. We observed INFLAREs in tissue sections and bulk RNAseq data of UC patients, who have an increased risk of CRC^72^. Bulk deconvolution of TCGA data suggested that INFLAREs are present in colon adenocarcinoma, particularly in MSI-high tumours. These results are consistent with recent identification of gastric metaplasia related gene expression (including *MUC5AC, TFF2, AQP5* along with reduced *CDX2*) in serrated polyps, which are pre-cancerous lesions associated with MSI high CRC^73^. In addition, there is a link between *TFF3* (elevated in metaplastic surface foveolar and INFLAREs) and CRC progression and survival^74^.

In conclusion, we present an integrated single cell atlas along the GI tract as a resource to study GI cell populations in health, development and disease. Using our atlas, we identify and interrogate gastric mucous/pyloric gland metaplasia, leading to a molecular model for the origin and role of metaplastic cells in intestinal inflammation, with implications for therapies and progression from inflammation to cancer.

## Methods

### Patient samples and tissue processing

#### Healthy tissue from adults

Healthy adult gastrointestinal tissue was obtained by the Cambridge Biorepository of Translational Medicine (CBTM) from deceased transplant organ donors (n = 2) after ethical approval (REC 15/EE/0152, East of England - Cambridge South Research Ethics Committee) and informed consent from the donor families. Details of the GI regions processed and donor information are compiled in Supplementary Table 4. Donors were perfused with cold University of Wisconsin (UW) solution and fresh tissue was collected from the distal stomach (antrum/pylorus), duodenum and terminal ileum within 1h of circulatory arrest, and stored in HypoThermosol FRS preservation solution (Sigma, H4416) at 4°C until processing. Intestinal tissue was open longitudinally and rinsed with D-PBS and then processed to single-cell suspensions following standard protocols ^3,75^. For tissues from donor A68/759B (D105) epithelium and lamina propria (LP) were separated into different fractions by dissection. Epithelial cells were removed by washing the intestinal mucosa twice in Hank’s Balanced Salt Solution (HBSS) medium (Sigma-Aldrich) containing 5 mM EDTA (Thermo Fisher, 15575020), 10 mM HEPES (Gibco, 42401042), 2% (v/v) FCS supplemented with 10 mM ROCK inhibitor (Y-27632 (Merck, Y0503)) while shaking at 4°C for 20 minutes. Epithelial wash-offs were centrifuged at 300 g for 7 min at 4°C and incubated at 37°C with TrypLE (Thermo Fisher) supplemented with 0.1 mg/mL DNase I (Sigma, 11284932001) for 5 minutes. Cells were pelleted and filtered through a 40 μm cell strainer and resuspended in Advanced DMEM F12 (ThermoFisher, 12634028) with 10% (v/v) FCS. The remaining epithelium-depleted tissue was minced and incubated in digestion media (HBSS medium, 0.25 mg/mL Liberase TL (Roche, 5401020001) and 0.1 mg/mL DNase I (Sigma, 11284932001)) on a shaker at 37°C for up to 45 min. The tissue was gently homogenised using a P1000 pipette every 15 mins. For tissues from donor A68/770C (D99), full thickness tissue was diced with a scalpel and digested in digestion media, as described above. Cells were pelleted and filtered through a 70 μm strainer before proceeding to Chromium 10x Genomics single cell 5’ v2 protocol as per manufacturer’s instructions. Libraries were prepared according to manufacturer’s protocol and sequenced on an Illumina NovaSeq 6000 S2 flow cell with 50 bp paired-end reads.

#### Control tissue from preterm infants

Uninvolved tissue from preterm infants, between 23 and 31 post conception weeks (pcw), with necrotising enterocolitis (NEC), focal intestinal perforation or intestinal fistula (n = 4) were collected at the Neonatal Department of Newcastle upon Tyne Hospitals NHS Foundation Trust with consent and ethical approval as part of the SERVIS study (REC 10/H0908/39). Tissue was resected from the infant and placed immediately into ice cold PBS. Within 3 hours, samples were enzymatically dissociated into a single cell suspension using Collagenase Type IV (Worthington) for 30 minutes at 37°C. Cells were filtered with 100 µm cell strainer, treated with red blood cell lysis and filtered through a 35 µm strainer. Cells were stained with DAPI before FACS sorting, selecting only for live, single cells. Sorted cells were then loaded onto the Chromium Controller (10x Genomics) using the Single Cell Immune Profiling kits and subsequently sequenced as per the manufacturer’s protocol.

#### Disease tissue from Crohn’s disease, Ulcerative Colitis and Celiac disease patients

CD tissue used for validations was obtained from multiple sites. Adult CD surgical resections were collected from patients in the IBSEN III (Inflammatory Bowel Disease in South Eastern Norway) at Oslo University Hospital (n = 4, Oslo, Norway) or Hospital Clinic Barcelona (n = 9, Barcelona, Spain), and biopsy material was collected from patients undergoing colonoscopy at Addenbrookes Hospital Cambridge (n = 4, Cambridge, United Kingdom); all patients gave informed written consent. Fresh tissue was fixed in formalin and embedded in paraffin for subsequent immunostaining. Ulcerative Colitis tissue was also collected from Hospital Clinic Barcelona (n = 3) during colonic resections, with the same consent and tissue processing procedure. Celiac disease tissue was obtained from Oslo University hospital (n = 2) or the Oxford University Hospitals NHS Foundation Trust (OUHFT) celiac disease clinic (n = 2 treated celiac, n = 3 untreated celiac).

Duodenal biopsies from Oslo University hospital were collected from newly diagnosed untreated patients with celiac disease (n = 2), and subsequently fixed in formalin and embedded in paraffin for immunostaining. Mucosal pinch biopsies from the second part of the duodenum from OUHFT were obtained during gastroscopy of untreated celiac patients (n = 3) and treated celiac patients on a gluten-free diet (n = 2). Equivalent healthy control samples from OUHFT (n = 3) were obtained from patients undergoing gastroscopy with gastrointestinal symptoms without celiac disease. Biopsies were stored in MACS tissue storage solution (Miltenyi Biotec) before cryopreservation in freezing medium (Cryostor Cs10, Sigma-Aldrich). Samples were later recovered by thawing in a 37°C water bath and washed in 20 mL R10 (90% RPMI (Sigma-Aldrich), 10% FBS) before tissue dissociation. Epithelial cells were isolated using v1.11 of the published protocol (dx.doi.org/10.17504/protocols.io.bcb6isre) ^76^. After isolation, epithelial cells proceeded to single cell sequencing (10x Genomics Next GEM 5’ v1.1) as per the manufacturer’s protocol. Details of samples and metadata are available in Supplementary Table 4.

#### Ethical approval for collection of disease tissue

Tissue collected at Oslo University Hospital was approved by the Regional Committee for Medical Research Ethics Ethics (REK 20521/6544, REK 2015/946, and REK 2018/703, Health Region South-East, Norway) and comply with the Declaration of Helsinki. Tissue collected at Hospital Clinic Barcelona was approved by the Ethics Committee of Hospital Clinic Barcelona (HCB/2016/0389). Tissue from Addenbrookes Hospital was collected through the Addenbrookes - Human Research Tissue Bank HTA research licence no: 12315 (Cambridge University Hospitals Trust). Tissue collected at OUHFT was collected under the Oxford Gastrointestinal Illnesses Biobank (REC 21/TH/0206).

#### Single-molecule fluorescence in situ hybridization

Intestinal tissue was embedded in OCT and frozen on an isopentane-dry ice slurry at −60 °C, and then cryosectioned onto SuperFrost Plus slides at a thickness of 10 μm. Before staining, tissue sections were post-fixed in 4% paraformaldehyde in PBS for 15 min at 4 °C, then dehydrated through a series of 50%, 70%, 100% and 100% ethanol, for 5 min each. Staining with the RNAscope Multiplex Fluorescent Reagent Kit v2 Assay (Advanced Cell Diagnostics, Bio-Techne) was automated using a Leica BOND RX, according to the manufacturers’ instructions. After manual pre-treatment, automated processing included epitope retrieval by protease digestion with Protease IV for 30 min prior to RNAscope probe hybridization and channel development with Opal 520, Opal 570, and Opal 650 dyes (Akoya Biosciences). Stained sections were imaged with a Perkin Elmer Opera Phenix High-Content Screening System, in confocal mode with 1 μm z-step size, using a 20x water-immersion objective (NA 0.16, 0.299 μm per pixel). Channels: DAPI (excitation 375 nm, emission 435–480 nm), Opal 520 (ex. 488 nm, em. 500–550 nm), Opal 570 (ex. 561 nm, em. 570–630 nm), Opal 650 (ex. 640 nm, em. 650–760 nm). The fourth channel was developed using TSA-biotin (TSA Plus Biotin Kit, Perkin Elmer) and streptavidin-conjugated Atto 425 (Sigma-Aldrich).

#### Immunohistochemistry

For samples collected at Oslo University hospital: Sections of FFPE tissue were cut in series at 4 µm and mounted on Superfrost Plus object glasses (ThermoFischer Scientific). Haematoxylin-eosin (H&E) staining was performed on the first sections and reviewed by an expert pathologist (FLJ) and the following sections were used for immunohistochemical studies. AB-PAS staining was performed by dewaxing FFPE samples and staining with Alcian blue (8GX) (AB) at pH 2.5 for acidic mucins and periodic acid-Schiff reagent (PAS) staining for neutral mucins, as described previously ^77^.

Multiplex immunostaining was performed sequentially using a Ventana Discovery Ultra automated slide stainer (Ventana Medical System, Roche, Cat. No. 750-601). After deparaffinization of the sections, heat-induced epitope retrieval was performed by boiling the sections for 48 minutes with Cell Conditioning 1 buffer (DISC CC1 RUO, Roche, 6414575001) followed by incubation with DISC inhibitor (Roche, 7017944001) for 8 minutes. The following primary antibodies were used: anti-human MUC6 clone CLH5 dilution 1:400. (RA0224-C.1, Scytek), anti-human MUC5AC clone CLH2 dilution 1:100 (MAB2011, Sigma), anti-human CD3 rabbit polyclonal dilution 1:50 (A0452, Dako), anti-human CD8 clone 4B11 dilution 1:30 (MA1-80231, Leica Biosystems, Invitrogen), anti-human CD4 clone SP35 dilution 1:30 (MA5-16338, Thermo Fisher), anti-TCR delta clone H-41 dilution 1:100 (sc-100289, Santa Cruz Biotechnology), Foxp3 clone 236A/E7 dilution 1:1000 (NBP-43316, Novus Biologicals), HLA-DR alpha-chain clone TAL.1B5 dilution 1:200 (M0746, Dako), CD68 clone PG-M1 dilution 1:100 (M0876, Dako), CD20 clone L26 dilution 1:200 (M0755, Dako).

Each primary antibody was diluted in Antibody Diluent (Roche, 5266319001), incubated for 32 minutes, and then washed in a 1x reaction buffer (Concentrate (10X), Roche, 5353955001). OmniMap anti-Mouse HRP (Roche, 5269652001) secondary antibody was incubated for 16 minutes followed by 12 minutes incubation with diluted opal fluorophores (Opal 6-Plex Detection Kit - for Whole Slide Imaging formerly Opal Polaris 7 Color IHC Automated Detection Kit NEL871001KT) following the manufacturer instructions. After that, bound antibodies were denatured and HRP was quenched using Ribo CC solution (DISC. CC2, Roche, 5266297001) and DISC. Inhibitor (Roche, 7017944001). Sections were then counterstained with DAPI (DISC QD DAPI RUO, Roche, 5268826001) for 8 min and mounted with ProLong Glass Antifade mountant (Molecular probes). Imaging was performed using a Vectra Polaris multispectral whole slide scanner (PerkinElmer). Irrelevant, concentration-matched primary antibodies were used as negative controls. For some tissue sections, bound anti-CD3, CD20, MUC6 and MUC5AC primary antibodies were detected with secondary antibodies conjugated with peroxidase, using the automated Ventana Discovery Ultra system and DAB, Purple, Yellow or Teal -responsive chromogens (ChromoMap DAB Detection Kit, 5266645001; DISCOVERY Purple Kit, 07053983001; DISCOVERY Yellow Kit, 07698445001; and Discovery Teal-HRP detection kit) all from Ventana Medical System.

For samples collected at Hospital Clinic Barcelona: Sections of FFPE tissue were cut into 3.5µm sections. IHC was conducted for the following commercially available antibodies: anti-mouse MUC5AC (Sigma-Aldrich, MAB2011, 1:4000) and anti-mouse MUC6 (ScyTek, RA0224-C.1, 1:4000). Deparaffinization, rehydration and epitope retrieval of the sections was automatedly performed with PT link (Agilent, CA, USA) using Envision Flex Target Retrieval Solution Low pH (Dako, Germany). Samples were blocked with 20% of goat serum (Vector, NY, USA) in a PBS and 0.5% BSA solution. Biotinylated anti-mouse secondary antibodies were used (1:200, Vector, NY, USA). Positivity was detected with the DAB Substrate kit (K3468, Dako, Germany). Image acquisition was performed on a Nikon Ti microscope (Japan) using Nis-Elements Basic Research Software (version 5.30.05).

#### Data curation and mapping

Datasets (Supplementary Table 1) were chosen from literature search of scRNAseq studies. Studies were included when there was raw scRNAseq data (FASTQ) from human GI tract tissues (oral cavity (excluding tongue), salivary glands, oesophagus, stomach, small and large intestine).

For public datasets deposited to ArrayExpress, archived paired-end FASTQ files were downloaded from ENA or ArrayExpress. For public datasets deposited to GEO, if the SRA archive did not contain the barcode read, URLs for the submitted 10X BAM files were obtained using srapath v2.11.0. The bam files were then downloaded and converted to fastq files using 10x bamtofastq v1.3.2. If the SRA archive did contain the barcode read, SRA archives were downloaded from the ENA and converted to FASTQ files using fastq-dump v2.11.0. Sample metadata was gathered from the abstracts deposited to GEO or ArrayExpress, and supplementary files from publications.

Following the FASTQ file generation, 10X Chromium scRNA-seq experiments were processed using the STARsolo pipeline v1.0 detailed in https://github.com/cellgeni/STARsolo repository. Briefly, STAR v2.7.9a was used. Transcriptome reference exactly matching Cell Ranger 2020-A for human was prepared as described in the 10X online protocol: https://support.10xgenomics.com/single-cell-gene-expression/software/release-notes/build#header. Automated script “starsolo_10x_auto.sh” was used to automatically infer sample type (3’ or 5’, 10X kit version, etc). STARsolo command optimised to generate the results maximally similar to Cell Ranger v6 was used. To this end, the following parameters were used to specify UMI collapsing, barcode collapsing, and read clipping algorithms: “-- soloUMIdedup 1MM_CR --soloCBmatchWLtype 1MM_multi_Nbase_pseudocounts -- soloUMIfiltering MultiGeneUMI_CR --clipAdapterType CellRanger4 --outFilterScoreMin 30”. For cell filtering, the EmptyDrops algorithm employed in Cell Ranger v4 and above was invoked using “--soloCellFilter EmptyDrops_CR” options. Options “--soloFeatures Gene GeneFull Velocyto” were used to generate both exon-only and full length (pre-mRNA) gene counts, as well as RNA velocity output matrices.

Following read alignment and quantification, Cellbender v0.2.0 with default parameters was used to remove ambient RNA (soup). In cases where the model learning curve did not indicate convergence, the script was re-run with “--learning-rate 0.00005 --epochs 300” parameters. For certain large datasets or datasets with low UMI counts, “--expected-cells” and “--low-count-threshold” parameters had to be adjusted individually for each sample.

#### Automated quality control

On a per sample basis, scAutoQC calculated the following metrics: number of counts and genes, percentage of ribosomal, mitochondrial and haemoglobin genes, and technical factors such as percentage of top 50 highly expressed genes, RNA soup and spliced genes (Extended data 2). The dimensions of these 8 metrics were reduced to generate a neighbourhood graph and UMAP, which was then clustered at low resolution - these clusters are referred to as QC clusters. Classification of cells/droplets as passing or failing QC was then performed in a two step process, firstly by classifying each cell as passing or failing QC based on 4 metric parameters and thresholds set by a Gaussian Mixture Model. Then, whole clusters were classified as passing QC if ≥ 50% of individual cells within the cluster passed QC. The benefits of the approach include the automated nature, removing most manually set thresholds and limiting hands-on analysis. Our unbiased approach exploits both the distribution of individual metrics and their correlations. Although there are some parameters which are set up-front, they only serve as guidance for the final flagging of low quality cells and are not sensitive to small changes in the starting points (for example, setting an initial % mitochondrial genes to 15 or 20% is likely to flag the same clusters). An overview of the pipeline is in Extended data 2 and the code (https://github.com/Teichlab/sctk/blob/master/sctk/_pipeline.py v0.1.1) and example workflow (https://teichlab.github.io/sctk/index.html) can be found in GitHub .

#### Assembly of the healthy reference

After samples were run through scAutoQC, they were pooled and cells flagged as failing QC, along with samples where < 10% of cells or 100 cells total passed QC (18 samples). Cells were further filtered through doublet removal (both automated, Extended data 2, and manual removal during annotations, see below). Cells from healthy/control samples were integrated using scVI ^78,79^ with donorID_unified as batch key, log1p_n_counts and percent_mito as continuous covariates, cell cycle genes removed and 7500 highly variable genes.

#### Annotations of the healthy reference

Cells from the core atlas were grouped by scanpy leiden clustering into broad categories of Epithelial, T and NK, B and B plasma, Myeloid, Mesenchymal, Neural and Endothelial based on marker gene expression (annotation level 1). Each lineage was split, and reintegrated with scVI (scVI settings) to annotate cells at fine resolution (annotation level 3). Mesenchymal populations were further split by developmental age group (first trimester fetal, second trimester fetal/preterm and adult/pediatric). Epithelial cells were further split by GI region and/or developmental age group (oral all ages, oesophagus all ages, stomach all ages, small intestine first trimester fetal, small intestine second trimester fetal/preterm, small intestine adult/pediatric, large intestine first trimester, large intestine second trimester fetal/preterm, large intestine adult/pediatric). For fine-grained annotations of objects by broad compartment (and age/region if applicable), a combined approach including automated annotation with leiden clustering and marker gene analysis was used. Celltypist predicted labels were calculated for the entire core atlas using various relevant models (Cells_Intestinal_Tract v2 - Elmentaite 2021, Immune_All_Low v2 - Dominguez Conde 2022 and Pan_Fetal_Human v2 - Suo 2022) and custom label transfer models from Martin 2019 (intestinal dataset), Warner 2020 (salivary gland dataset). Notebooks for all annotations are available via our GitHub. MGN cells (MUC6+) in the healthy reference in small intestine were identified in healthy duodenum with leiden clustering resolution 0.5, and further refined to remove any residual doublets or MUC6-cells by subclustering.

#### Disease data projection and label prediction

Models for disease projection were made on the full healthy reference dataset (without doublets) using scANVI incorporating broad (level 1) annotations, based on the healthy reference scVI model. We projected disease data using scArches with the scANVI model. To annotate at fine resolution, we first predicted broad (level 1) lineages in the projected disease data using a label transfer method based on majority voting from kNN. Broad lineages were then split as for the healthy reference. For all lineages except epithelial, lineage specific disease cells were projected onto the respective healthy reference lineage specific latent space and fine-grained annotations predicted using the same method as for broad lineage predictions. Due to an underrepresentation of epithelial cells, we added additional epithelial cell data from celiac disease duodenum (unpublished data from Klenerman lab - Fitzpatrick2023) and CD ileum and colon ^24^, increasing the amount of diseased epithelial cells from 57,406 to 92,342 cells plus an additional 219,472 cells from healthy controls/non-inflammed tissue. These additional datasets were not remapped, instead these studies were added based on the raw counts matrix. Split epithelial cells from the original disease set (remapped data) and the additional disease sets (from count matrices) were concatenated and reduced to a common gene set of 18,485 genes. The resulting epithelial dataset was further split by region (stomach, small intestine and large intestine), prepared for projection using scANVI_prepare_anndata function (fills 0s for non-overlapping genes) and projected onto the respective healthy reference epithelial region specific latent space embeddings.

To refine level 3 annotations on disease cells, we utilised scArches weighted kNN uncertainty metric. We labelled cells as unknown if they had an ‘uncertainty score’ greater than the 90th quantile for each lineage. For epithelial cells the 90th quantile was calculated separately for cells from cancer and non-cancer to account for high uncertainty labelling of tumour cells. To refine the labels of these unknown cells, we performed leiden clustering (resolution = 1) and reassigned the label based on both majority voting of the higher certainty cells (above the cut off) and marker genes. In stomach epithelium, there was one cluster of unknown cells, likely to be cancer cells, which could not be assigned a label and was therefore left annotated as unknown. In large intestine epithelium we found a cluster which corresponded to metaplastic Paneth cells (a cell type not present in the healthy reference) which were reannotated based on the distinct marker genes (Extended data 4).

#### Quantitation of technical and biological variation

To determine the contribution of different metadata covariates to the integrated embedding of the healthy reference data, we performed linear regression for each latent component of the embedding with each covariate as previously described^59^. We performed the analysis per cell type based on level_1_annot (broad level) and level_2_annot (medium level) annotations, and for all ages or adult/pediatric only (excluding developing and preterm samples).

#### Differential abundance analysis

To identify differentially abundant cell populations, we used Milo^31^ which tests for differentially abundant neighbourhoods from kNN graphs. For comparisons between healthy developing (6-31 pcw, including preterm infants ex utero) and adult/pediatric gut, Milo was run separately per tissue with > 2 donors for each group (stomach, duodenum, ileum and colon) using default parameters. For comparisons between organs in healthy adult gut (≥ 18 years), Milo was run for each organ (oral mucosa, salivary gland, oesophagus, stomach, small intestine, large intestine, mesenteric lymph node) vs the others combined with the covariates of tissue_fraction and cell_fraction_unified and otherwise default parameters. For comparisons between disease and healthy adult samples, Milo was run comparing disease and controls from an individual study, rather than all disease and controls in the atlas, on the kNN graph from joint embedding which has been shown to have greater sensitivity for detecting disease associated cell states ^32^. We focused comparison inflamed with neighbouring inflamed tissue from Martin19 dataset,

#### Differential gene expression, gene set analysis and gene scoring

Differential gene expression analysis was performed by Wilcoxon rank sum (Mann-Whitney U) test using Scanpy rank gene groups function with default parameters. To limit unwanted batch and technical effects, samples were preprocessed by downsampling to 200 cells per cell type per donor and removing ribosomal and mitochondrial related genes. For Gene set analysis, the output from Scanpy rank gene groups was filtered to contain genes with a minimum log fold change of 0.25 and p-value cut off of 0.05. The resulting gene list was used for Gene set analysis using GSEApy enrichr function with relevant genesets such as MSigDB, KEGG and GO_Biological_Process examined. Gene scores for epithelial cells were calculated using Drug2Cell^80^ score function with default parameters. Gene scores for fibroblasts were calculated using Scanpy score_genes function with default parameters. Full gene lists used for gene scores are available in Supplementary Table 5.

#### Cell-cell interaction analysis

Cell-cell interaction analysis was performed using CellChat^81^ and CellPhoneDB v4 (statistical_method) ^82^ to determine cell-cell interactions occurring in the small intestine during CD. Interaction analysis was performed on remapped data, to avoid loss of genes/interactions lost when merging additional count matrices (See Disease data projection and label prediction for more detail). Before analysis, data was preprocessed by downsampling to 50 cells per cell type per donor. Normalised count matrix with cell annotation metadata were processed through the standard CellChat and CellPhoneDB pipeline, with the communication probability truncated mean/threshold set to 0.1.

#### cNMF analysis

To identify shared activity and cell identity gene programs cells from diseased small intestine (CD, pediatric IBD and celiac disease with a total of 99,465 cells), we analysed raw counts with consensus non-negative matrix factorization (cNMF) 1.3.4^83^. We used the default processing and normalisation of cNMF, which considers 2,000 highly variable genes along with 100 iterations of NMF. All other parameters were set at default values. We tested hyperparameter values of K, the number of factors, ranging in steps of 1 from 5-80 and picked on inspection a favourable tradeoff between factor stability and overall model error at K=44. For determining consensus clusters we excluded 6% of fitted cNMF spectra with a mean distance to k-nearest neighbours above 0.3. The resulting per-cell gene program usage was compared across fine-grained cell annotations, identifying gene programs corresponding to the identity of Mucous_gland_neck cells and other relevant cell types (Goblet, Stem and Surface foveolar cells).

#### Trajectory analysis

To infer the developmental trajectory giving rise to MGN/INFLARE cells in the ileum IBD we used monocle3 ^84^ on a subset of data containing cells in the ileum from Jaeger2021 (GSE157477), Martin2019 (GSE134809) and Kong2023 (SCP1884). We performed Louvain clustering on the original UMAP representation generated from the scANVI latent space to account for batch effects and inferred developmental trajectories along pseudotime by choosing the node assigned the highest number of epithelial stem cells as the root node. We then extracted the MGN/INFLARE-specific trajectory by selecting the nodes assigned the highest number of MGN/INFLARE cells as the final nodes. Finally, we determined genes whose expression changes along pseudotime by using ‘monocle3::graph_test’, which leverages a Moran’s I test considering gene expression changes within groups of k=25 neighbouring cells on the principle trajectory graph.

#### Bulk RNA sequencing deconvolution using single cell data

For bulk deconvolution analysis, we first downloaded published bulk RNA-seq datasets of adult IBD from the Gene Expression Omnibus (GEO) database (GSE111889), pediatric IBD from the ArrayExpress database (E-MTAB-5464) and the Expression Atlas (E-GEOD-101794), TCGA colon adenocarcinoma using R package TCGAbiolinks and celiac disease data from GEO (GSE131705 and GSE145358). A single-cell reference for deconvolution analysis was then prepared by subsetting the overall object to only include cells from the small intestine in disease conditions and downsampling to 300 cells for each fine-grained cell type annotation. BayesPrism^43^ was used for deconvolution analysis with raw counts for both single-cell and bulk RNA-seq data as inputs. Both the ‘cell type labels’ and the ‘cell state labels’ were set to fine-grained annotations. Ribosomal protein genes and mitochondrial genes were removed from single-cell data as they are not informative in distinguishing cell types and can be a source of large spurious variance. We also excluded genes from sex chromosomes and lowly transcribed as recommended by the BayesPrism tutorial. For further analysis, we applied a pairwise t-test to select differentially expressed genes with the ‘pval.max’ being set to 0.05 and ‘lfc.min’ to 0.1. Finally, a prism object containing all data required for running BayesPrism was created using the new.prism() function, and the deconvolution was performed using the run.prism() function. For correlation analysis, we calculated the Pearson correlation between (1) the estimated abundance of INFLAREs and other cell types, and (2) the estimated INFLAREs abundance and gene expression in bulk RNA-seq datasets. For the later calculation, we first normalised raw counts in the expression matrix from each bulk dataset using R package DESeq2.

### Data Availability statement

All unpublished transcriptomic data will be available upon or within 6 months of publication. Published datasets are readily available through public repositories, single cell datasets included in the atlas are listed in Supplementary Table 2. Accession numbers for published bulk RNAseq datasets are included in the methods section. Imaging data will be available for download from the European Bioinformatics Institute (EBI) BioImage Archive upon publication. All relevant processed single cell objects and models for use in future projects will be available upon publication, through gutcellatlas.org.

### Code Availability statement

Code for scAutoQC is readily available on GitHub and installable via PyPI (https://github.com/Teichlab/sctk). Additional code including atlas assembly, annotation and downstream analyses is described in detail throughout the methods, and will be available on GitHub upon publication.

## Acknowledgements

We thank the donors and their families for donating tissue samples and enabling this research. We also thank the organisers and members of the Helmsley Consortium and Human Cell Atlas Gut Bionetwork for facilitating valuable discussions. We thank K. Roberts for support and expertise with imaging; A. Oszlanczi for her help on sample management; A. Wilk for administrative assistance; H. H. Uhlig and M. G. Friedrich for valuable discussions, L. Buer and K. Thorvaldsen Hagen for assistance with tissue blocks and AB-PAS staining. We acknowledge support from the Wellcome Sanger Cellular Genetics Informatics team, Spatial Genomics Platform (SGP) team, particularly M. Patel, Core DNA Pipelines and New Pipeline Group (NPG), especially S. Leonard. We are grateful to the Cambridge Biorepository for Translational Medicine for access to tissue from deceased transplant organ donors. This work was made possible through collaboration between the Wellcome Sanger Institute, University of Oslo and Oslo University Hospital, IDIBAPS Hospital Clinic Barcelona, Newcastle University, Cambridge University Hospitals NHS Foundation Trust, University of Cambridge and the University of Oxford. This work was financially supported by the Wellcome Trust (WT206194, S.A.T.); the European Research Council (646794, ThDefine, S.A.T.); an MRC New Investigator Research Grant (MR/T001917/1, M.Z.); and a project grant from the Great Ormond Street Hospital Children’s Charity, Sparks (V4519, M.Z.). A.M.C and V.G. were funded by grant Grant #2008-04050 from The Leona and Harry B. Helmsley Charitable Trust. E.M.A is funded by grant RH042155 (RTI2018-096946-B-I00) from Ministerio de Ciencia e Innovacion. This study was supported by NIHR Biomedical Research Centre, Oxford. The views expressed in this paper are those of the author/s and not necessarily those of the NHS, the NIHR or the Department of Health. The illustration in Figure 5f was made using BioRender https://biorender.com, all other illustrations were made by R.E and A.J.O. This publication is part of the Human Cell Atlas: https://www.humancellatlas.org/publications.

## Conflicts of interest

In the past 3 years, S.A.T. has consulted or been a member of scientific advisory boards at Roche, Genentech, Biogen, GlaxoSmithKline, Qiagen and ForeSite Labs and is an equity holder of Transition Bio and EnsoCell. R.E. is an equity holder in EnsoCell. P.K. has consulted for AstraZeneca, UCB, Biomunex and Infinitopes. J.S-R reports funding from GSK, Pfizer and Sanofi and fees/honoraria from Travere Therapeutics, Stadapharm, Astex, Pfizer and Grunenthal.

## Author information

**Project design:** A.J.O., R.E., S.A.T. **Data generation (tissue procurement):** M.E.B.F, T. R.W.O., C.E.H., K.T.M., K.S-P., M.L.H., E.S.B., A.S., C.S., J.B. **Data generation (tissue processing and sequencing):** R.B-C., N.M.P., J.C., A.M., E.S., J.E., L.R., R.K., A.W-C., C.I.S., A.G-T., R.E. **Data generation (RNAscope):** S.E., C.T., P.J. **Data generation (IHC and histology):** H.N., V.G., E.M-A., S.P. **Data curation:** A.J.O., R.L., R.B-C., R.E. **Data analysis (raw data processing):** N.H., A.V.P., B.C., **Data analysis (QC):** N.H. **Data analysis (atlas assembly):** N.H., A.J.O. **Data analysis (atlas annotation):** A.J.O., R.L., R.B-C., R.E. **Data analysis (disease mapping and annotation):** A.J.O. **Data analysis (healthy analysis):** A.J.O., R.L., K.P. **Data analysis (disease analysis):** A.J.O., R.L., S.K., L.M.M., B.C., K.T., R.E. **Data analysis (tissue validation):** A.J.O., R.B-C., R.E. **Data analysis (bulk deconvolution):** R.L. **scAutoQC package:** N.H., K.P., S.L. **Gut Cell Survey web portal:** M.P. **Project insight:** A.J.O., R.L., R.B-C., E.D., J.S-R., K.R.J., K.B.M., M.Z., A.S., P.K., M.H., F.L.J., R.E., S.A.T. **Review and editing:** A.J.O., R.L., R.B-C., S.L., M.M., R.E., S.A.T. **Writing:** A.J.O., R.E., S.A.T. **Supervision:** A.W-C., J.S-R., K.B.M., A.S., P.K., M.H., F.L.J., S.A.T.

## Extended data

**Extended data 1:**
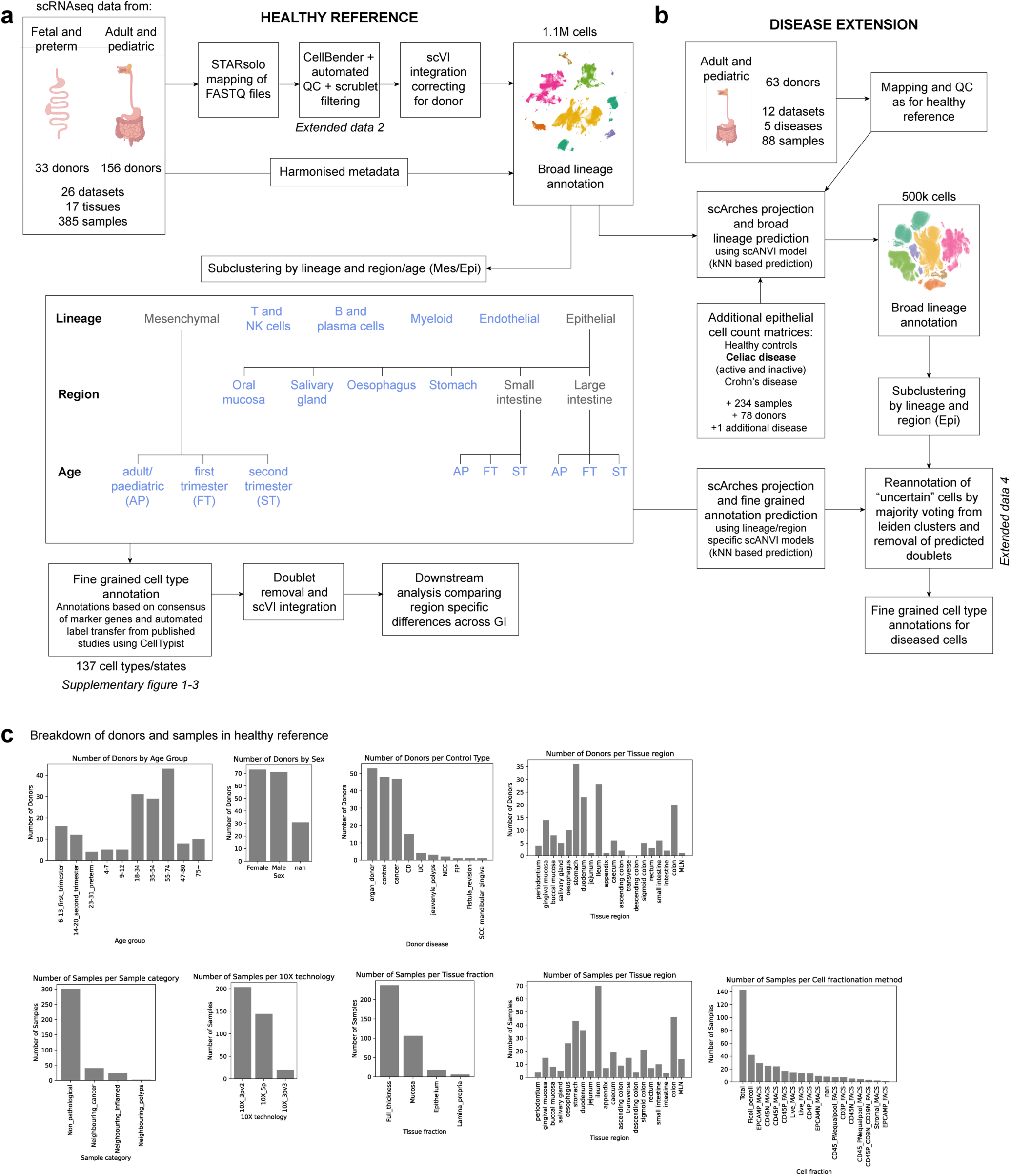
Overview of atlas assembly. a) Detailed flowchart of the methods used to assemble the healthy reference, datasets were remapped and filtered based on scAutoQC automated QC pipeline (Supplementary Figure 2), integrated with scVI and annotated as broad lineages. Broad lineages were subclustered, and lineages with high level of heterogeneity (Epithelial and Mesenchymal lineages) were further subclustered based on age and/or region to accurately annotate at a fine-grained level. Cells in these subclustered views of the healthy reference were annotated by a semi-automated approach, taking into account the marker genes and CellTypist predictions from published studies. b) The healthy reference was used as an anchor to project disease datasets onto the atlas using scArches, fine-grained annotations were generated in a two-step approach, first with broad lineage prediction using scANVI and subclustering by lineage/region as with the healthy reference to predict the fine-grained annotations. Most disease data was remapped and QC’ed as with the healthy reference, except two additional studies from CD (Kong2023) and celiac disease (Fitzpatrick2023) which were added to the atlas from the raw count matrices. c) Breakdown of the distribution of donors and samples in the healthy reference based on metadata such as age, sex, disease, tissue sampling methods etc.

**Extended data 2:**
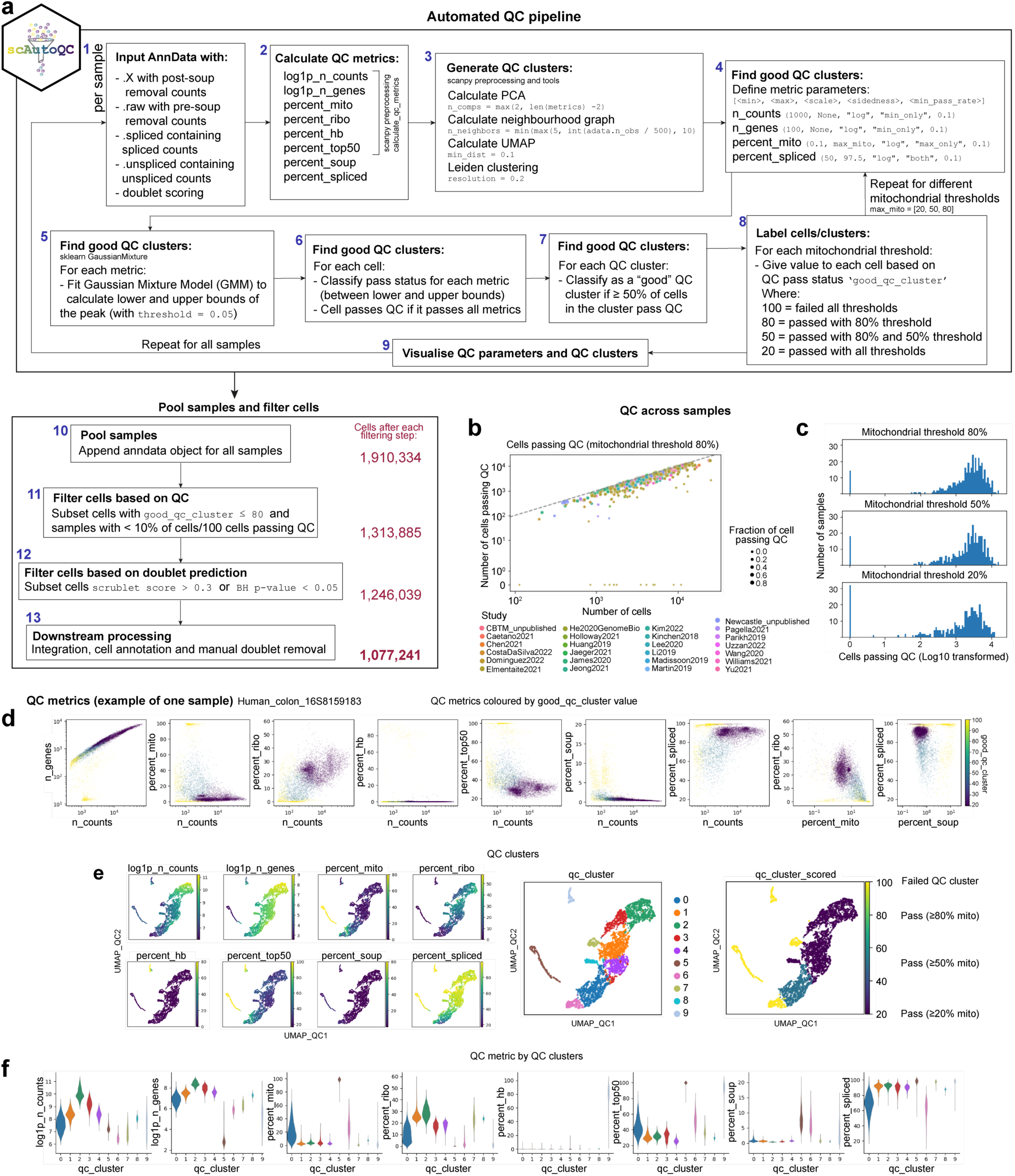
Overview of scAutoQC method. a) Summary of the automated QC pipeline. Standard QC metrics are calculated and dimensions of 8 QC metrics (listed in step 2) are reduced, neighbours calculated and UMAP generated. Clusters from this UMAP are classified as “good” if ≥ 50% fall within upper and lower bounds (calculated by Gaussian Mixture Model) of 4 QC metrics (listed in step 4). Step 4-7 was repeated for 3 different mitochondrial thresholds (20%, 50%, 80%) and all steps were repeated for all samples. Finally samples are pooled, and cells within clusters that failed automated QC when mitochondrial threshold is 80%, and predicted as doublets (based on scrublet score calculated on a per sample basis) are removed before downstream processing. b) Plot of cells passing QC vs number of cells per sample across studies. Dotted line represents threshold for 100% of cells/sample passing QC. c) Histogram showing distribution of cells passing QC (log base 10) across the 3 mitochondrial thresholds. d-f) Example QC plots from one sample where d) is showing QC distribution of QC metrics where each data point is a cell, coloured by good_qc_cluster value (see step 8 of panel a). e) shows the QC UMAPs with the 8 QC metrics (listed in step 2 panel a), QC leiden clusters and good_qc_cluster value (see step 8 of panel a). f) violin plot of the 8 QC metrics (listed in step 2 of panel a) for each QC leiden cluster. In this sample for example, cluster 5 has failed QC because cells in this cluster have high % of mitochondrial reads, low genes and high percentage of genes expressed within the top 50 genes.

**Extended data 3:**
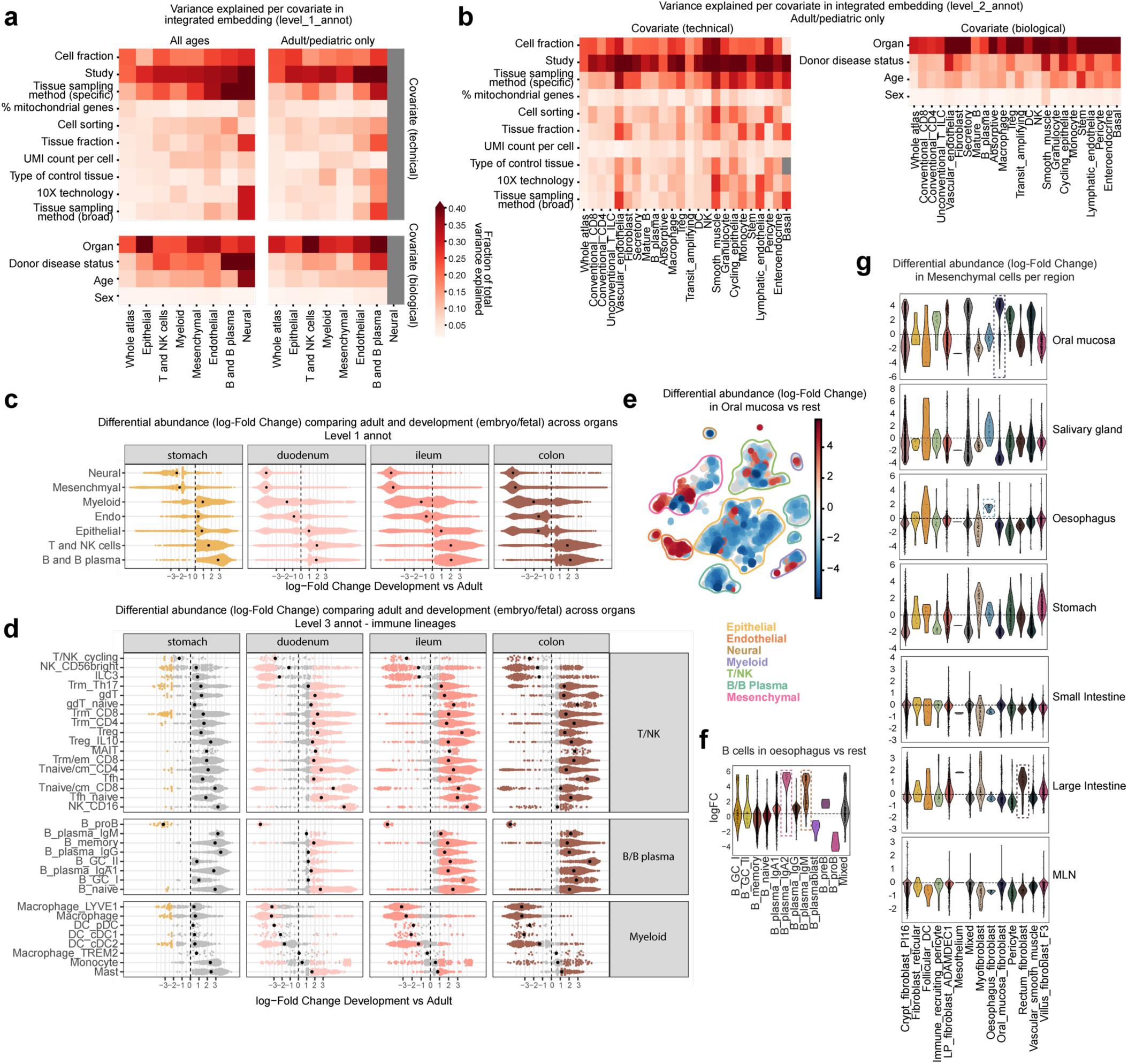
Analysis of cells within the healthy reference. a) Analysis of metadata covariate contribution of variance in the integrated healthy reference embedding per cell type at broad level annotations (level_1_annot). b) Analysis of covariate contribution of variance per cell type at mid-level annotations (level_2_annot). c) Differential abundance analysis (Milopy) comparing broad level cell type (level_1_annot) abundance between adult/pediatric samples and developing samples (embryo, fetal and preterm), broken down by GI region with sufficient data for comparison. Each datapoint is a neighbourhood with positive log-fold change values indicating enrichment of lineage in adult/pediatric GI vs developing GI. d) Differential abundance analysis (Milopy) comparing fine-grained cell type/state (level_3_annot) abundance from immune lineages between adult/pediatric samples and developing samples (embryo, fetal and preterm), broken down by GI region. Each datapoint is a neighbourhood with positive log-fold change values indicating enrichment of cell type/state in adult/pediatric GI vs developing GI. Coloured data points are significantly enriched/depleted neighbourhoods. e) UMAP showing differential abundant neighbourhoods in the healthy reference comparing Oral mucosa to other organs throughout the GI tract in adult/pediatric samples. Positive log-fold change indicates enrichment of neighbourhoods in Oral mucosa. Coloured neighbourhoods show significant enrichment/depletion. f) Violin plot of B and B plasma cells in oesophagus showing enrichment of IgA2 and IgM plasma cells. g) Differential abundance of Mesenchymal populations in adult/pediatric samples across each GI region compared to all others combined. Three tissue specific fibroblast populations were annotated, Oral mucosa, Oesophagus and Rectum fibroblasts.

**Extended data 4:**
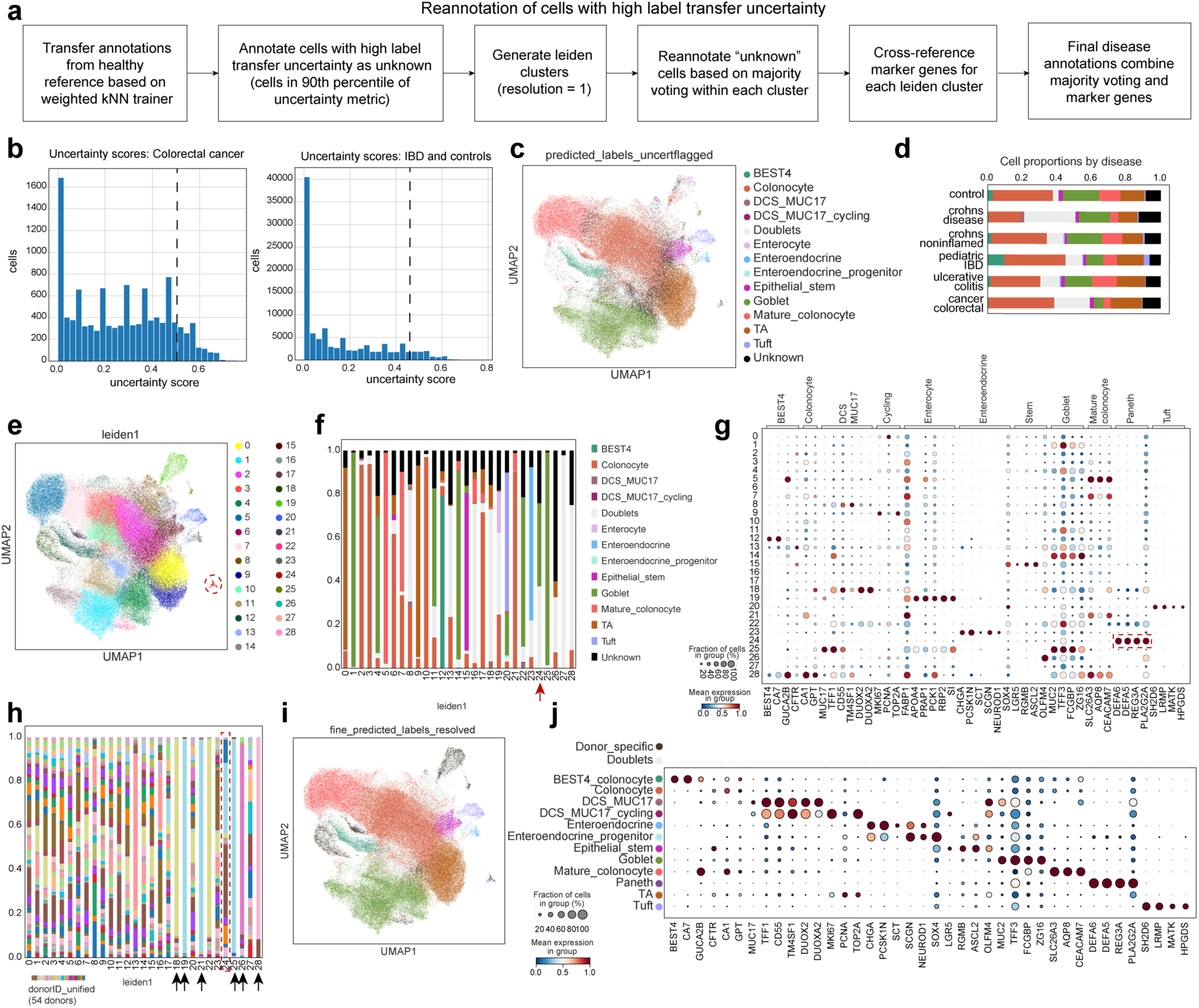
Workflow for annotation of disease data and identification of metaplastic Paneth cells. a) Flowchart for the annotation of cells from disease samples. As described in Extended data 1, cells from disease samples were added to the atlas by scArches projection. Annotations from the healthy reference were transferred by a weighted kNN classifier, cells with an uncertainty score within the 90th percentile were classified as unknown and reannotated based on majority voting of leiden clusters. All disease annotations were cross-referenced with marker gene expression from literature and through differential gene expression analysis. b-j) Example workflow for large intestine epithelial cells in disease. b) Distribution of uncertainty scores in disease data from large intestine epithelial cells from cancer and non-cancer. Dashed line indicates the 90th percentile cut off, where cells with an uncertainty score above this are classified as “unknown”. c) UMAP of large intestine epithelial cells with predicted annotations and unknown cells flagged. DCS = deep crypt secretory cells. d) Proportions of predicted large intestine epithelial cell annotations (colours as in c) including unknown cells by disease. e) UMAP of large intestine epithelial cells with leiden clustering at resolution = 1, used to reclassify unknown cells based on majority voting. f) Proportions of predicted large intestine epithelial cell annotations by leiden cluster. Red arrow points to cluster 24, which was reannotated to Paneth cells but originally annotated as a combination of goblet cells, doublets and unknown. g) Marker gene dot plot of large intestine epithelial cells and Paneth cells by leiden cluster. Paneth cell markers are highlighted for cluster 24. h) Proportions of cells in each leiden by donor. Black arrows highlight clusters dominated by cells from only one donor (later excluded from the atlas), and red arrow highlights cluster 24 which contains metaplastic Paneth cells. i) UMAP of reannotated large intestine epithelial cells from disease, including metaplastic Paneth cells. j) Marker gene dot plot for reannotated cell types in large intestine epithelial cells from disease.

**Extended data 5:**
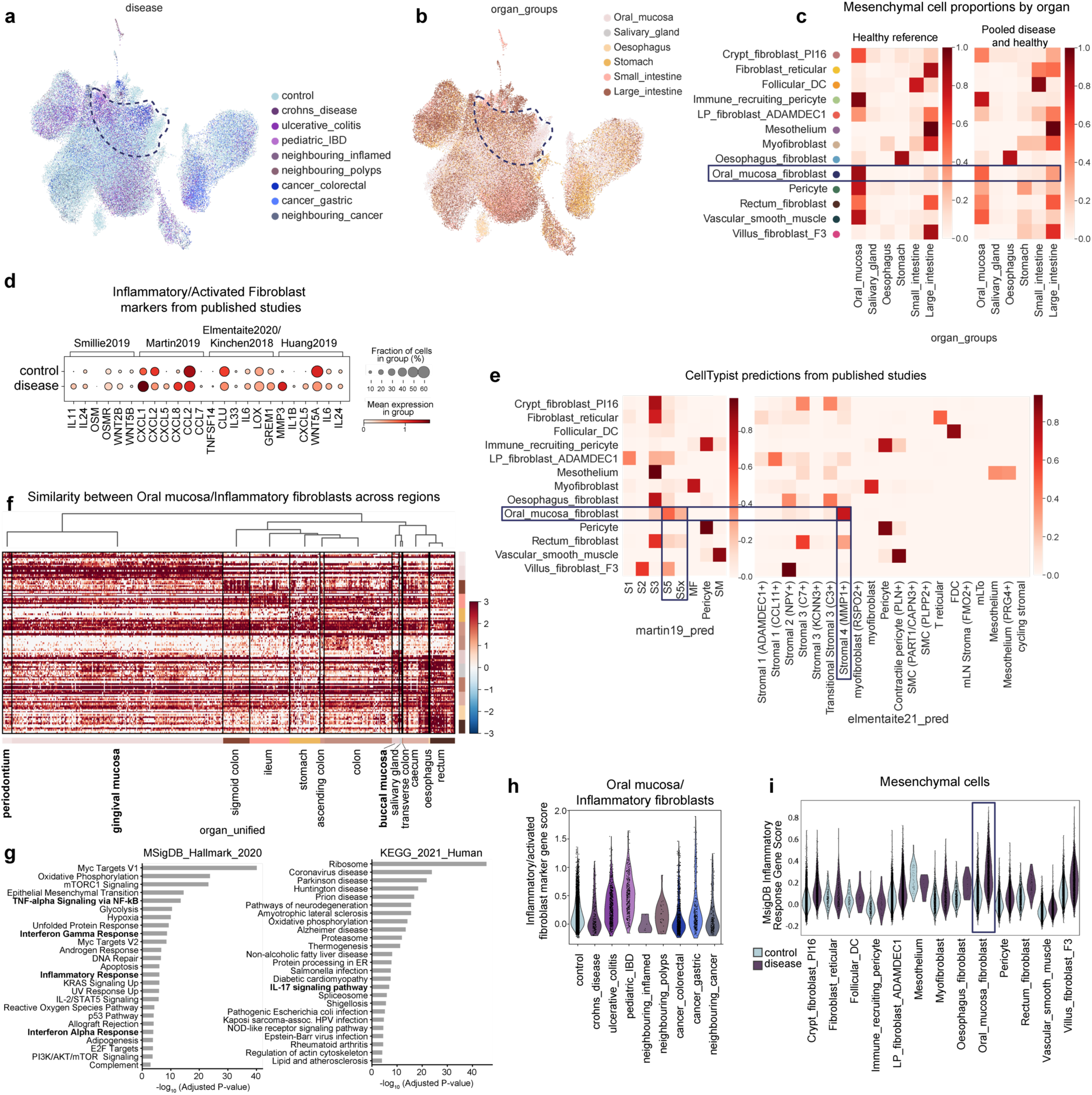
Inflammatory fibroblasts in disease are annotated as Oral mucosa fibroblasts. a) UMAP of mesenchymal cells from adult/pediatric samples in health and disease, shown by disease category. Dashed line highlights the oral mucosa fibroblast cluster. b) UMAP of mesenchymal cells from adult/pediatric samples in health and disease, shown by organ. Dashed line highlights the oral mucosa fibroblast cluster. c) Proportion of mesenchymal cell types/states by organ in the healthy reference and combined healthy and disease. Oral mucosa fibroblasts appear in other organs in disease. d) Markers of inflammatory and activated fibroblasts from published studies^27^ showing expression in oral mucosa/inflammatory fibroblasts from controls (oral mucosa fibroblasts) and disease (inflammatory fibroblasts) samples. e) CellTypist predictions of cell annotations in mesenchymal populations from published studies ^3,4^ showing oral mucosa fibroblasts predicted to be inflammatory/activated fibroblast populations in both studies. f) Differential gene expression (wilcoxon rank sum test) and hierarchical clustering of oral mucosa/Inflammatory fibroblasts from different regions. Oral mucosa fibroblasts from gingival mucosa and periodontium are most distinct from fibroblasts in other organs. g) Gene set enrichment analysis showing pathways (including various inflammatory pathways) enriched in inflammatory fibroblasts (disease) compared to oral mucosa fibroblasts (healthy). h) Gene score for inflammatory/activated fibroblasts markers in (e) expressed in oral mucosa/inflammatory fibroblasts across disease conditions. i) MSigDB inflammatory response gene score (significantly enriched in inflammatory vs oral mucosa fibroblasts), across all mesenchymal cell types/states in control and disease samples.

**Extended data 6:**
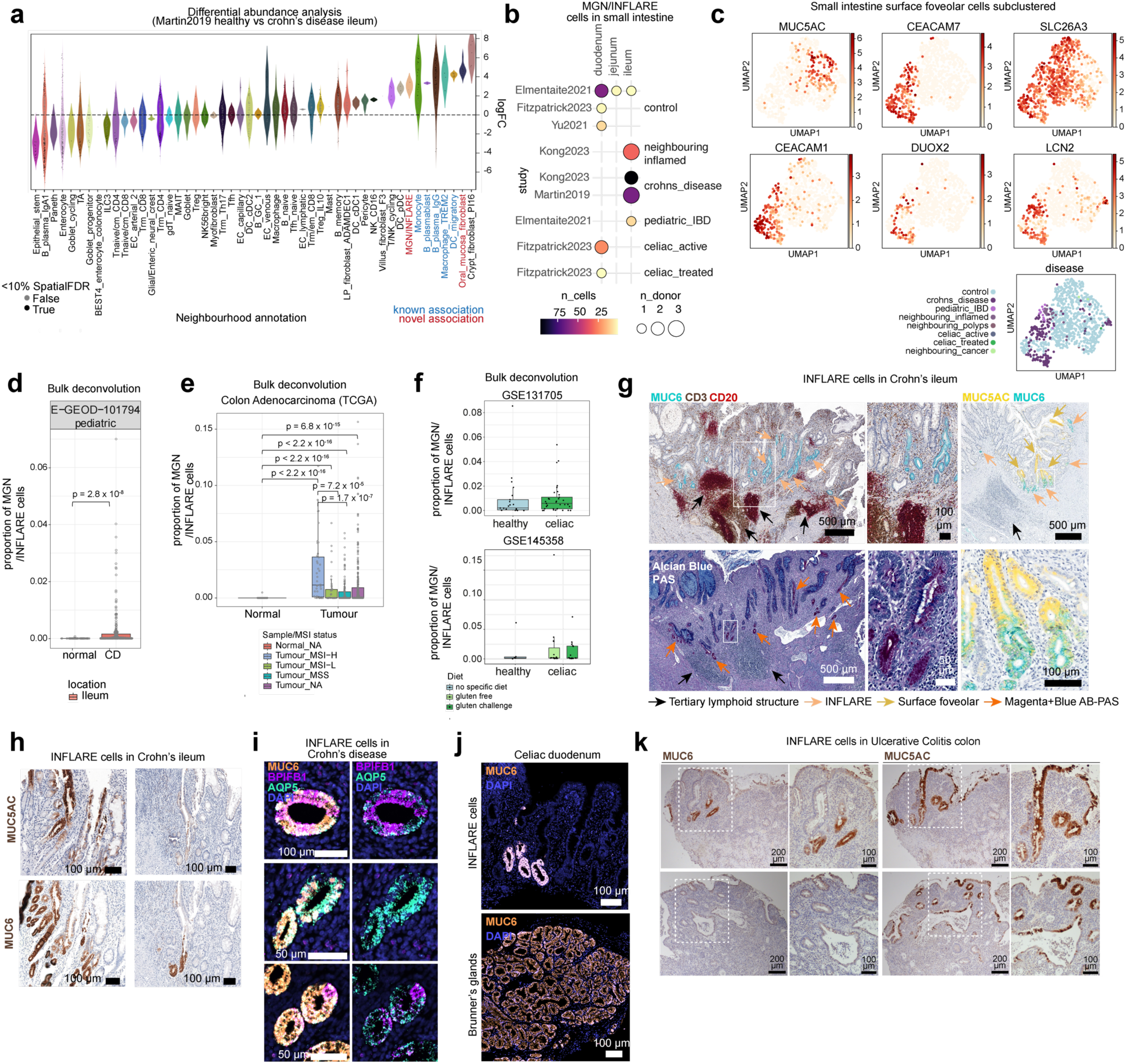
Validation of INFLARE cells. a) Differential abundance analysis of cell neighbourhoods from Martin 2019 dataset based on embedding on the whole atlas. Cell neighbourhoods with positive log fold change are enriched in CD compared to healthy samples. b) Overview of the number of MGN (Mucous gland neck)/INFLAREs (<u>Infla</u>mmatory <u>E</u>pithelial cell<u>s</u>) and donors per study, broken down by age and region of the GI. Dot size indicates the number of donors, colour indicates the number of cells. c) Subclustering of surface foveolar cells, showing heterogeneity of marker genes and additional genes upregulated in metaplastic surface foveolar cells. d) Deconvolution (BayesPrism) of bulk RNAseq dataset comparing MGN/INFLAREs in healthy (normal, n = 50) and CD (n = 254). e) Deconvolution (BayesPrism) of TCGA bulk RNAseq data of MGN/INFLAREs in healthy tissue (normal, n = 41) and tumour tissue stratified by microinstability status, n = 40 (Tumour_MSI-H), n = 42 (Tumour_MSI-L), n = 126 (Tumour_MSS) and n = 272 (Tumour_NA). MSI-high tumours are predicted to have higher levels of INFLAREs. f) Deconvolution (BayesPrism) of bulk RNAseq from celiac disease comparing MGN/INFLARE proportions in healthy and celiac disease tissue. For GSE131705, n = 21 (healthy) and n = 33 (celiac). For GSE145358, n = 6 (healthy), n = 15 (celiac gluten free) and n = 15 (celiac gluten challenge). For all box and whisker plots the lower edge, upper edge and centre of the box represent the 25th (Q1) percentile, 75th (Q3) percentile and the median, respectively. The interquartile range (IQR) is Q3 - Q1. Outliers are values beyond the whiskers (upper, Q3 + 1.5 x IQR; lower, Q1 - 1.5 x IQR). g) Protein and ABPAS staining of INFLAREs (MUC6, Magenta+Blue+ ABPAS staining) and Surface foveolar cells (MUC5AC) in CD ileum showing association with tertiary lymphoid structures (dense nuclei and CD3/CD20+ regions). h) Protein staining of INFLAREs (MUC6) and Surface foveolar cells (MUC5AC) from CD ileum tissue from additional donors. i) smFISH staining of INFLARE (<u>Infla</u>mmatory <u>E</u>pithelial cell) markers (*MUC6, AQP5* and *BPIFB1*) in pyloric metaplasia of CD duodenum showing heterogeneity in *AQP5* and *BPIFB1* expression. j) Protein staining of INFLAREs (MUC6) in celiac disease duodenum from two separate donors. k) Protein staining of INFLAREs (MUC6) and Surface foveolar cells (MUC5AC) in colon resection tissue from ulcerative colitis patients. Upper and lower panels are images from two different patients.

**Extended data 7:**
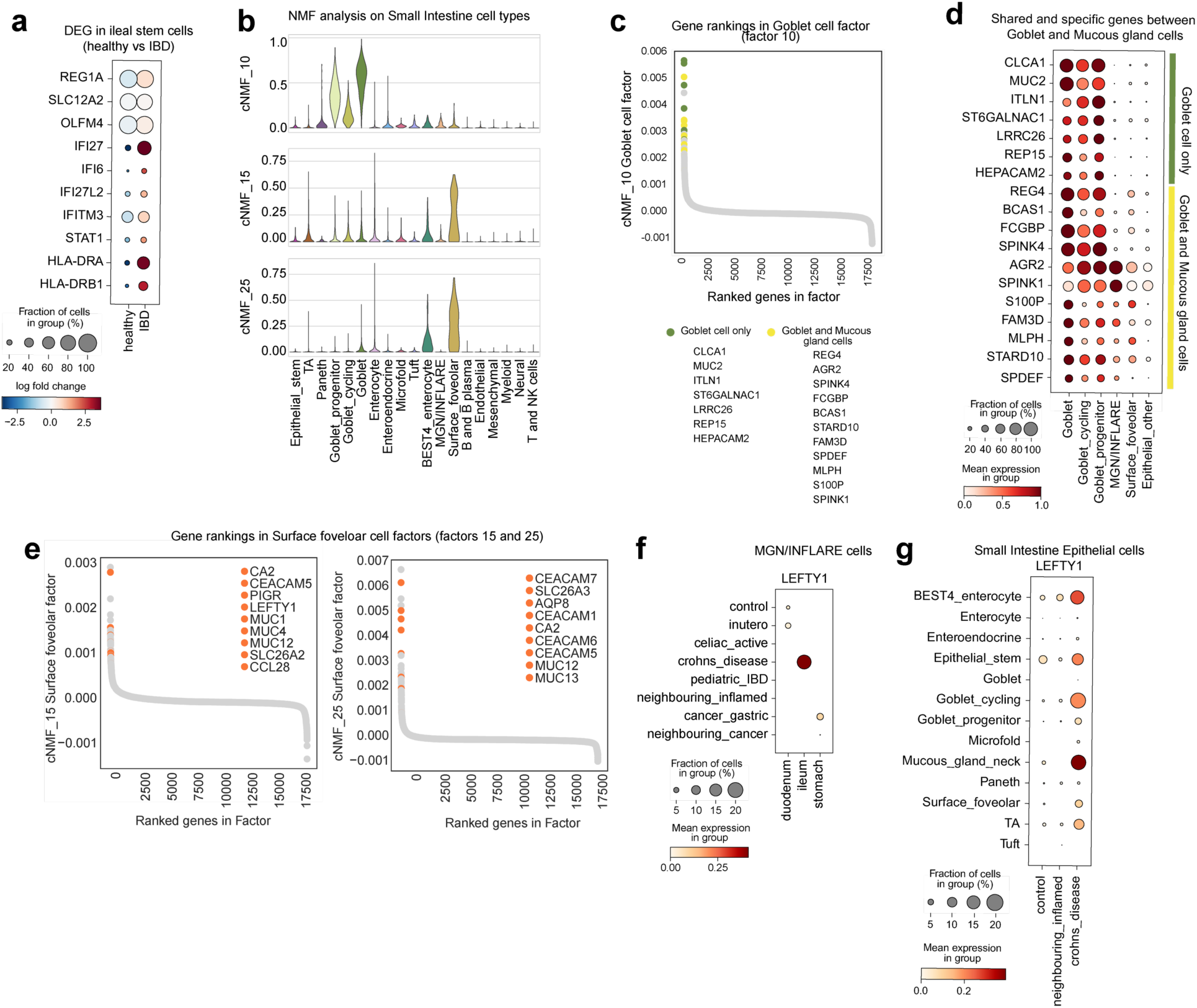
Origins and stem-like features of INFLAREs. a) Dot plot of selected significantly differential expressed genes in epithelial stem cells (*LGR5+*) from the ileum of patients with IBD compared with healthy controls. b) Non-negative factorisation analysis (Methods) of cell types from small intestine in the atlas. Violin plots showing expression of ranked genes in factors related to surface foveolar and goblet cells. c) Gene rankings of genes in factor 10 (goblet cell factor) with goblet cell specific genes highlighted in green and those also expressed in Mucous gland cells (MGN/INFLARE and surface foveolar cells) highlighted in yellow. d) Dotplot of genes highlighted in (c) showing expression in goblet cell populations, MGN/INFLAREs, Surface foveolar cells and all other epithelial cells in the small intestine. e) Gene rankings of genes in factors 15 and 25 (surface foveolar cell factors) with select genes highlighted. f) Dotplot of *LEFTY1* expression in MGN/INFLARE cells across regions and conditions. g) Dotplot of *LEFTY1* expression in all small intestinal epithelial cells, by cell annotation and condition. Crohn’s disease includes adult and pediatric Crohn’s disease.

**Extended data 8:**
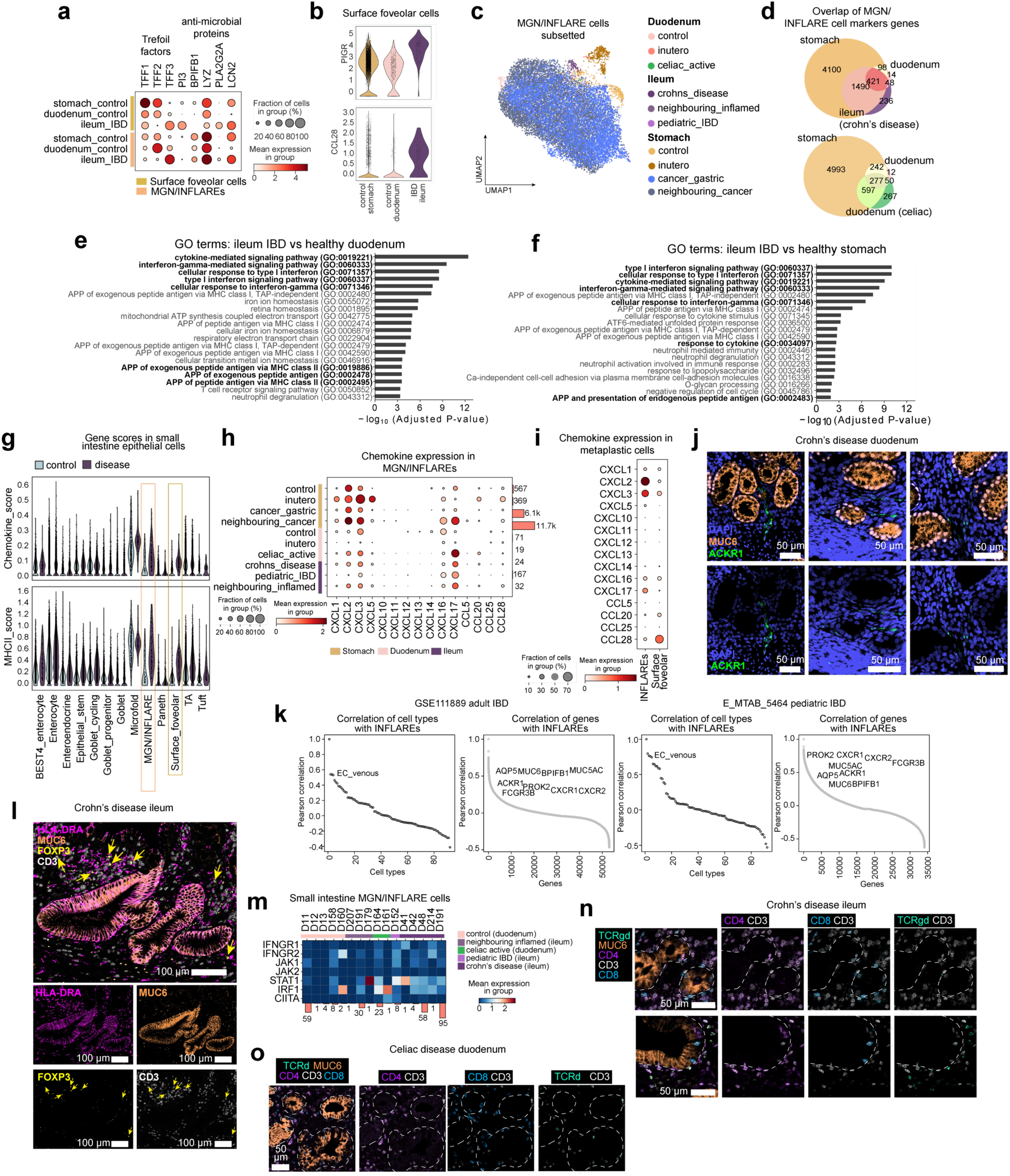
Dual role of gastric mucous metaplasia in mucosal healing and inflammation. a) Expression of genes related to mucosal barrier function in MGN (Mucous gland neck)/INFLAREs (<u>Infla</u>mmatory <u>E</u>pithelial cell<u>s</u>) and Surface foveolar cells in healthy stomach, healthy duodenum and IBD ileum. b) Violin plot showing expression of *PIGR* and *CCL28*, genes involved in mucosal IgA response in Surface foveolar cells from healthy stomach, healthy duodenum and IBD ileum. c) Subclustered MGN/INFLARE cells from across the atlas (locations, ages and diseases). MGN/INFLARE cells from different regions and duodenum/stomach in utero occupy separate coordinates in the UMAP. d) Overlap of MGN/INFLARE marker genes from different regions. Marker genes of MGN/INFLARE cells were calculated by differential gene expression (wilcoxon rank sum test) of other stomach and small intestine epithelial cells separately for healthy adult stomach MGN, healthy adult duodenum MGN, ileum CD INFLARE and duodenum celiac disease INFLARE. Overlapping marker genes show greater similarity of INFLAREs to healthy adult stomach MGN cells, than to healthy adult duodenum MGN cells. e) GO terms from upregulated genes (wilcoxon rank sum test) in IBD INFLARE cells (CD and pediatric IBD) compared with healthy control duodenum. Highlighted pathways are inflammatory, MHC-II mediated antigen presentation and exogenous peptide antigen presentation related pathways. f) Analysis as in (e) comparing IBD INFLARE cells to healthy control stomach. Comparison to healthy control stomach did not highlight MHC-II related pathways as in the comparison to healthy control duodenum. g) Chemokine and MHC-II gene scores (see Supplementary Table 5 for gene list) comparing small intestine epithelial cells in the atlas in healthy control and disease (IBD and celiac) samples showing specificity of upregulated chemokine and MHC-II related gene expression in metaplastic gastric mucous glands, particularly in INFLAREs vs MGN cells. h) Expression of chemokines expressed by MGN/INFLARE cells, across healthy and diseased tissues. i) Expression of chemokines on metaplastic INFLARE and surface foveolar cells, specifically in diseased small intestine (pooled cells from IBD and celiac disease). j) Additional smFISH staining (as in Figure 5c) of INFLAREs (*MUC6*) association with ACKR1+ vessel in CD duodenum. k) Correlation between INFLARE cell proportions and cell types/genes from deconvolution (BayesPrism) of bulk RNAseq adult and pediatric IBD datasets using the atlas as a reference. Analysis indicates consistent correlation of EC_venous cells (ACKR1+ endothelial population) with INFLAREs, and Surface foveolar and neutrophil marker genes with INFLAREs. l) Protein expression in CD ileum of HLA-DR (MHC-II) in INFLAREs (MUC6) along with localisation of CD3+ T cells and regulatory T cells (FoxP3+CD3+). m) Expression per donor of genes involved in IFNGR to MHC-II signalling pathway in INFLAREs and MGN cells in small intestine, as summarised in Figure 5f. n) Additional protein staining for INFLAREs (MUC6) in CD disease ileum (as in Figure 5g) with various T cell subsets (CD4+CD3+, CD8+CD3+, TCRγδ+CD3+ T cells). o) Protein staining as in (n) in Celiac disease duodenum tissue.

## Supplementary files

**Supplementary figure 1:**
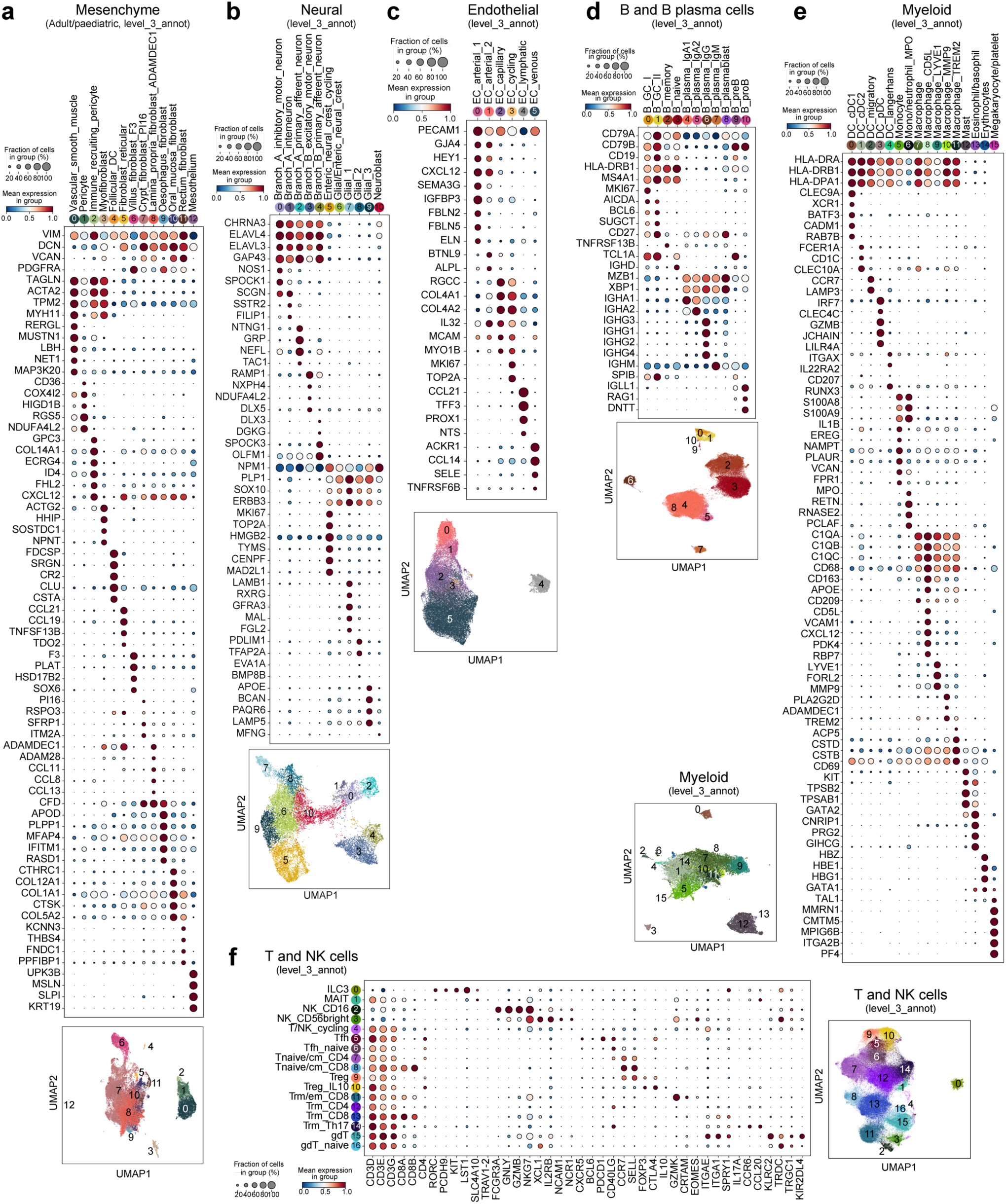
Fine-grained annotations with marker dot plot and UMAP of cells from non-epithelial lineages. a) Mesenchymal lineage annotations for cells in adult/pediatric samples. b) Neural lineage annotations. c) Endothelial lineage annotations. d) B and B plasma lineage annotations. e) Myeloid lineage annotations. f) T and NK lineage annotations.

**Supplementary figure 2:**
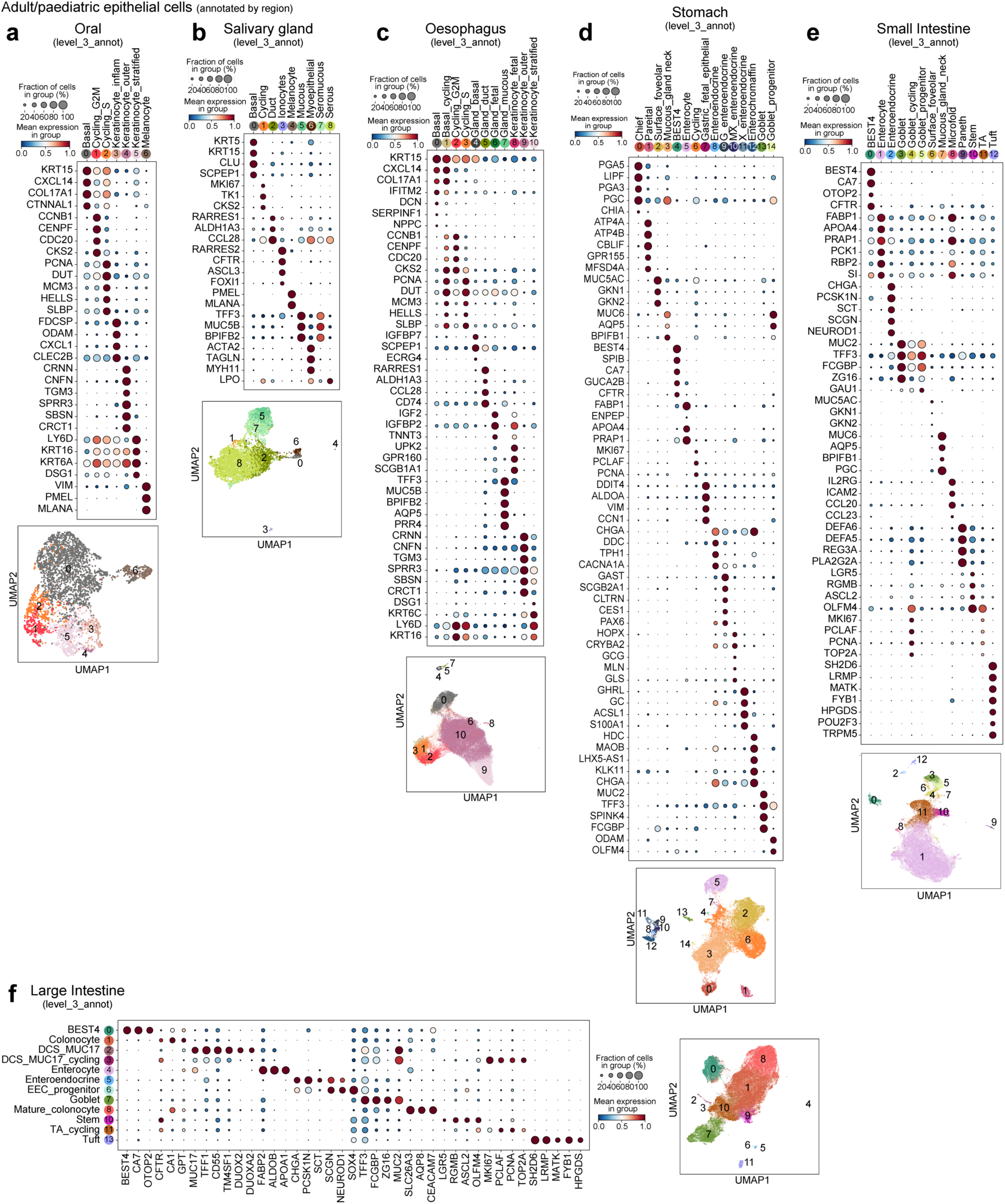
Fine-grained annotations with marker dot plot and UMAP of cells from adult/pediatric epithelial lineages, subclustered by organs. a) Oral mucosa epithelial cells (periodontium, gingival and buccal mucosa). b) Salivary gland epithelial cells. c) Oesophagus epithelial cells. d) Stomach epithelial cells. e) Small intestine epithelial cells (duodenum, jejunum, ileum). f) Large intestine epithelial cells (appendix, ceacum, ascending/descending/transverse/sigmoid colon, rectum).

**Supplementary figure 3:**
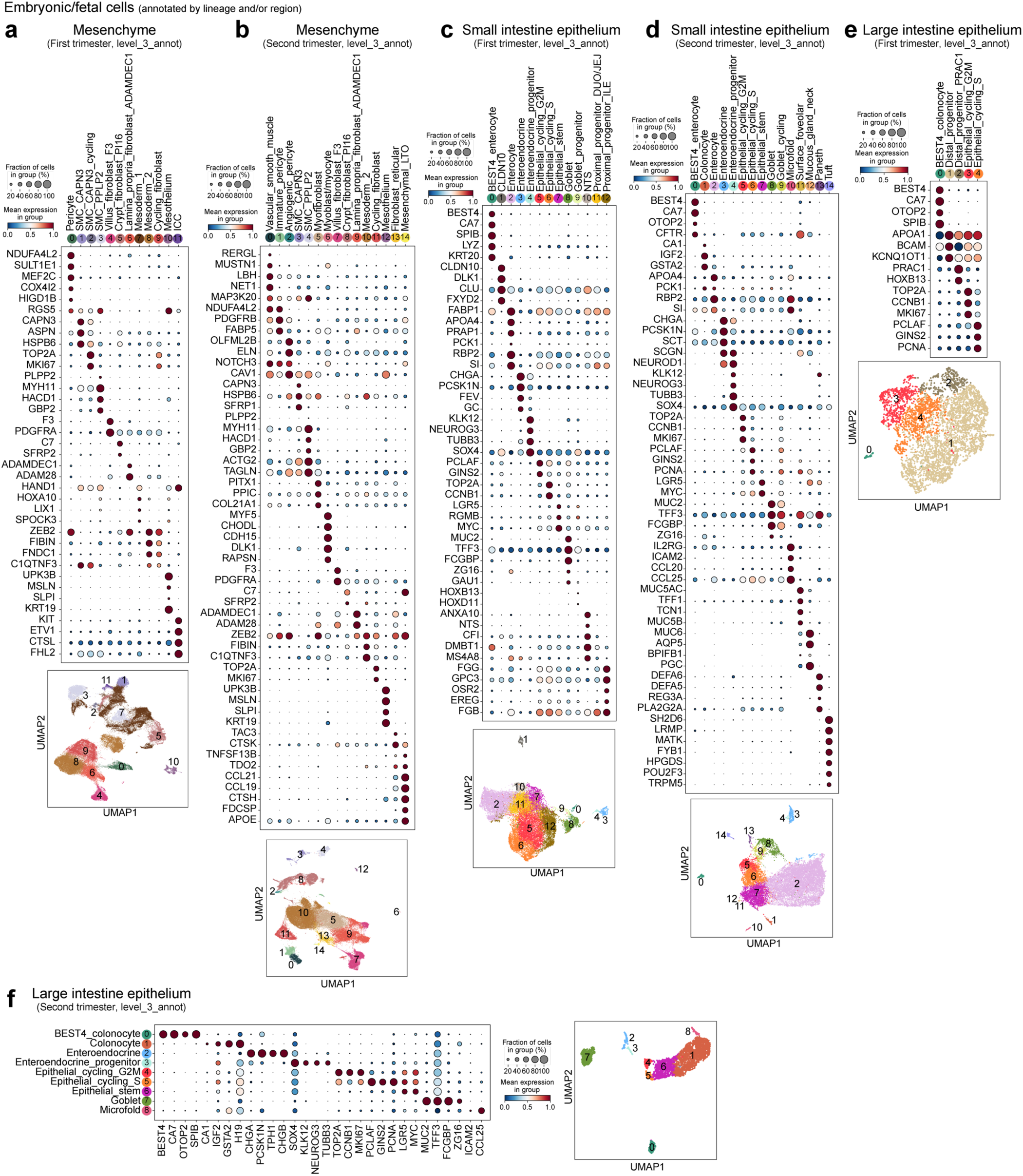
Fine-grained annotations with marker dot plot and UMAP of mesenchymal and small/large intestine epithelial cells from embryo, fetal and preterm samples. a) Mesenchymal cells from first trimester samples. b) Mesenchymal cells from second trimester second trimester and preterm samples. c) Small intestine epithelial cells from first trimester samples. d) Small intestine epithelial cell second trimester and preterm samples. e) Large intestine epithelial cells from first trimester samples. f) Large intestine epithelial cell second trimester and preterm samples.

## Supplementary tables

**Supplementary Table 1:** Overview of studies included in the atlas

**Supplementary Table 2:** Harmonized metadata included in the atlas

**Supplementary Table 3:** Overview of patient samples used for validation of INFLAREs

**Supplementary Table 4:** Overview of unpublished single cell data included in the atlas

**Supplementary Table 5:** Gene lists used for gene scoring

## Notes

https://www.gutcellatlas.org/

### Competing Interest Statement

S.A.T. is a scientific advisory board member of ForeSite Labs, OMass Therapeutics, a co-founder and equity holder of TransitionBio and EnsoCell Therapeutics, a non-executive director of 10x Genomics and a part-time employee of GlaxoSmithKline. R.E. is an equity holder in EnsoCell. P.K. has consulted for AstraZeneca, UCB, Biomunex and Infinitopes. N.M.P reports consulting fees from Infinitopes. J.S.-R. reports funding from GSK, Pfizer and Sanofi and fees/honoraria from Travere Therapeutics, Stadapharm, Astex, Owkin, Pfizer, Moderna and Grunenthal. A.S. is the recipient of research grants from Roche-Genentech, Abbvie, GSK, Scipher Medicine, Pfizer, Alimentiv, Boehringer Ingelheim and Agomab and has received consulting fees from Genentech, GSK, Pfizer, HotSpot Therapeutics, Alimentiv, Agomab, Goodgut and Orikine. R.E. and S.A.T. are inventors on the patent GB2412853.0 filed in the UK, some components of which are related to this work. All other authors declare no competing interests.

